# Investigating the Synergistic Role of GCN2 and HPA Axis in Regulating Integrated Stress Response in the Central Circadian Timing System

**DOI:** 10.1101/2024.02.01.578301

**Authors:** Yannuo Li, Lingjun Lu, Jordan L. Levy, Tracy G. Anthony, Ioannis P. Androulakis

## Abstract

The circadian timing and integrated stress response (ISR) systems are fundamental regulatory mechanisms that maintain body homeostasis. The central circadian pacemaker in the suprachiasmatic nucleus (SCN) governs daily rhythms through interactions with peripheral oscillators via the hypothalamus-pituitary-adrenal (HPA) axis. On the other hand, ISR signaling is pivotal for preserving cellular homeostasis in response to physiological changes. Notably, disrupted circadian rhythms are observed in cases of impaired ISR signaling. In this work, we examine the potential interplay between the central circadian system and the ISR, mainly through the SCN and HPA axis. We introduce a semi-mechanistic mathematical model to delineate the suprachiasmatic nucleus (SCN)’s capacity for indirectly perceiving physiological stress through glucocorticoid-mediated feedback from the HPA axis, and orchestrating a cellular response via the ISR mechanism. Key components of our investigation include evaluating general control nonderepressible 2 (GCN2) expression in the SCN, the effect of physiological stress stimuli on the HPA axis, and the interconnected feedback between the HPA and SCN. Simulation reveals a critical role for GCN2 in linking ISR with circadian rhythms. Notably, a *Gcn2* deletion in mice led to swift re-entrainment of the circadian clock post simulated-jetlag. This is attributed to the diminished robustness of neuronal oscillators and an extended circadian period. Our model also offers insights into phase shifts induced by acute physiological stress and the alignment/misalignment of physiological stress with external light-dark cues. Such understanding aids in strategizing responses to stressful events, such as nutritional status changes and jetlag.

## Introduction

The regulation of the central circadian timing system, mainly driven by the light-dark signal, is pivotal for upholding daily rhythms in physiology and behavior. Physiological stress, a state where the body perceives threats or challenges that disrupt homeostasis, has been shown to impact circadian rhythms (Koch et al. 2017; Pierre, Schlesinger, and Androulakis 2016). Recent developments have expanded our understanding of how physiological stress can affect the suprachiasmatic nucleus (SCN) – the central pacemaker of the circadian system – potentially altering its timing signals and downstream effects on the body. The extent to which physiological stress affects the central clock is debatable. Although some findings suggest the resilience of the SCN to unpredictable physiological stress stimuli due to the lack of glucocorticoid receptor (GR) expression (Tahara and Shibata 2018), there is evidence that acute or chronic physiological stressors can influence the central oscillator within the SCN. In cases of acute physiological stress, studies have shown that exogenous glucocorticoid surge can boost the expression of arginine vasopressin (AVP) and vasoactive intestinal peptide (VIP) mRNA within the SCN (Larsen et al. 1994). Chronic unpredictable stress (CUS) in rats has also unveiled diminished PER2 oscillations in SCN neurons (Jiang et al. 2011), implying potential disruptions in SCN rhythmicity from chronic physiological stress.

Physiological stress signals, often linked to metabolic inputs such as feeding/fasting cycles and dietary nutrient intake, elicit a multifaceted bodily response (Foteinou et al. 2009). Neuroendocrine hormones orchestrate physiological stress responses at the systemic level, while the integrated stress response (ISR) addresses stress within the cellular environment (Zänkert et al. 2019; Galluzzi, Yamazaki, and Kroemer 2018).

Specifically, the anterior piriform cortex (APC) is a critical site for detecting physiological stress, such as the deficiency of essential amino acids (EAA). Here, general control nonderepressible 2 (GCN2), the eIF2α kinase, becomes active when it binds to uncharged tRNA, initiating the cellular ISR (Hao et al. 2005; Maurin et al. 2005). APC projections to the hypothalamus ensure ISR signals are integrated into the neuroendocrine response (Harding et al. 2003). The hypothalamic-pituitary-adrenal (HPA) axis then triggers a hormonal response influencing mood, digestion, immune function, and energy balance (Zänkert et al. 2019). Corticotropin-releasing hormone (CRH), particularly, is essential for the detection of physiological stress and, during EAA deficiency – as evidenced in studies with mice on a leucine-deficient diet – leads to increased activity of the sympathetic nervous system (Anthony and Gietzen 2013). Consequently, the APC and hypothalamus work together to modulate physiological and cellular stress responses, facilitating adaption to metabolic changes. This coordinated process is deeply influenced by the body’s EAA nutritional status, ensuring effective metabolic adaption.

Beyond influencing the hypothalamus, the physiological stress signal associated with metabolic shifts, such as those between feeding and fasting also impact the SCN. For instance, a decline in SCN neural activity has been observed just before and during feeding times in daylight hours, typically periods of heightened neural activity (Dattolo et al. 2016). This indicates the SCN’s capability to process information stemming from metabolic changes. The influence of glucose on SCN neural activity has also been documented (Hall et al. 1997), underscoring the SCN’s potential role in tracking nutritional status. However, the underlying mechanisms are not yet fully understood.

Central to these interactions is GCN2’s role in nutrition and metabolic stress signaling at the cellular level. Given its ubiquity in mammalian tissues, especially in the brain, liver, and skeletal muscle (Hu and Guo 2020), GCN2’s influence is significant. As the primary ISR sensor, GCN2 substantially shapes circadian physiology. In the SCN, heightened GCN2 activity shortens the circadian period, whereas its reduction prolongs and even disrupts rhythmicity (Pathak et al. 2019). This positions GCN2 within the SCN as potentially vital in interlinking the central circadian clock with metabolic stress and nutrition cues. However, the exact dynamics through which GCN2 processes ISR within the SCN remain to be elucidated.

The intricate interplay between the central circadian clock and integrated stress response underscores the importance of their harmonious balance for overall homeostasis. We propose a hypothesis centered on the dynamic interplay between the central circadian physiology and the ISR system, with a particular focus on metabolic stress induced by EAA deficiency, using it as a representative model for the effects of low protein quality in the diet. The ISR kinase GCN2, located in the SCN - the central circadian pacemaker – is activated by metabolic stress signals. This activation leads to the phosphorylation of eIF2α by GCN2, subsequently impacting clock gene transcription through the activating transcription factor 4 (ATF4) protein, as highlighted in the study by Pathak et al. (Pathak et al. 2019). Concurrently, the HPA axis, a principal physiological stress responder, responds to metabolic stress signals with an increase in CRH production. This heightened CRH output has the potential to communicate back to the SCN through glucocorticoid secretion. Thus, while the HPA axis orchestrates the systemic response to metabolic stress, GCN2 within the SCN detects cellular stress and adjusts the circadian rhythm in response to nutritional signals. This dual mechanism underscores a complex interplay between the ISR and circadian regulation.

To elucidate the intertwined dynamics of the central circadian physiology and ISR, we introduce a refined semi-mechanistic mathematical model. The model delves into the impact of GCN2 expression within the SCN, the influence of ISR stimuli on the HPA axis, and the intricate feedback loop connecting the HPA with the SCN. Our primary objective is to investigate the synergistic role of GCN2 and the HPA axis in orchestrating the stress response within the central circadian framework. In line with experimental observations, our simulation results suggest that GCN2 plays a critical role in connecting the integrated stress response with circadian rhythms. Specifically, the knockout of GCN2 results in a rapid re-entrainment of the circadian clock after jetlag due to the reduced robustness of neuronal oscillators and a lengthening of the circadian period. Our model investigates the effects of both acute and circulating physiological stress, including how the latter aligns with external time, providing a simplified framework for understanding how the central circadian compartment senses and interacts with physiological stress signals. The proposed mechanisms effectively integrate the current knowledge on the SCN’s role in nutrition and metabolic stress detection, as well as the HPA axis’s response to physiological stress. Additionally, the model takes into account the modulatory effects of glucocorticoids on the SCN, advancing our comprehensive understanding of the intricate equilibrium that underpins bodily homeostasis. This understanding contributes to the development of interventions to alleviate or assist in recovering from stressful events such as EAA deficiency and jetlag.

## Materials and Methods

The “**Model Development**” section in the **Supplementary Materials** provides a detailed explanation of the mathematical model formulated to elucidate the interplay between the central circadian clock and the ISR sensing mechanism. As depicted in **Figure 1**, the model captures the intricate interaction between the SCN and the HPA axis, with the former influenced by light signals and the latter by physiological stress signals.

**Figure 1.**
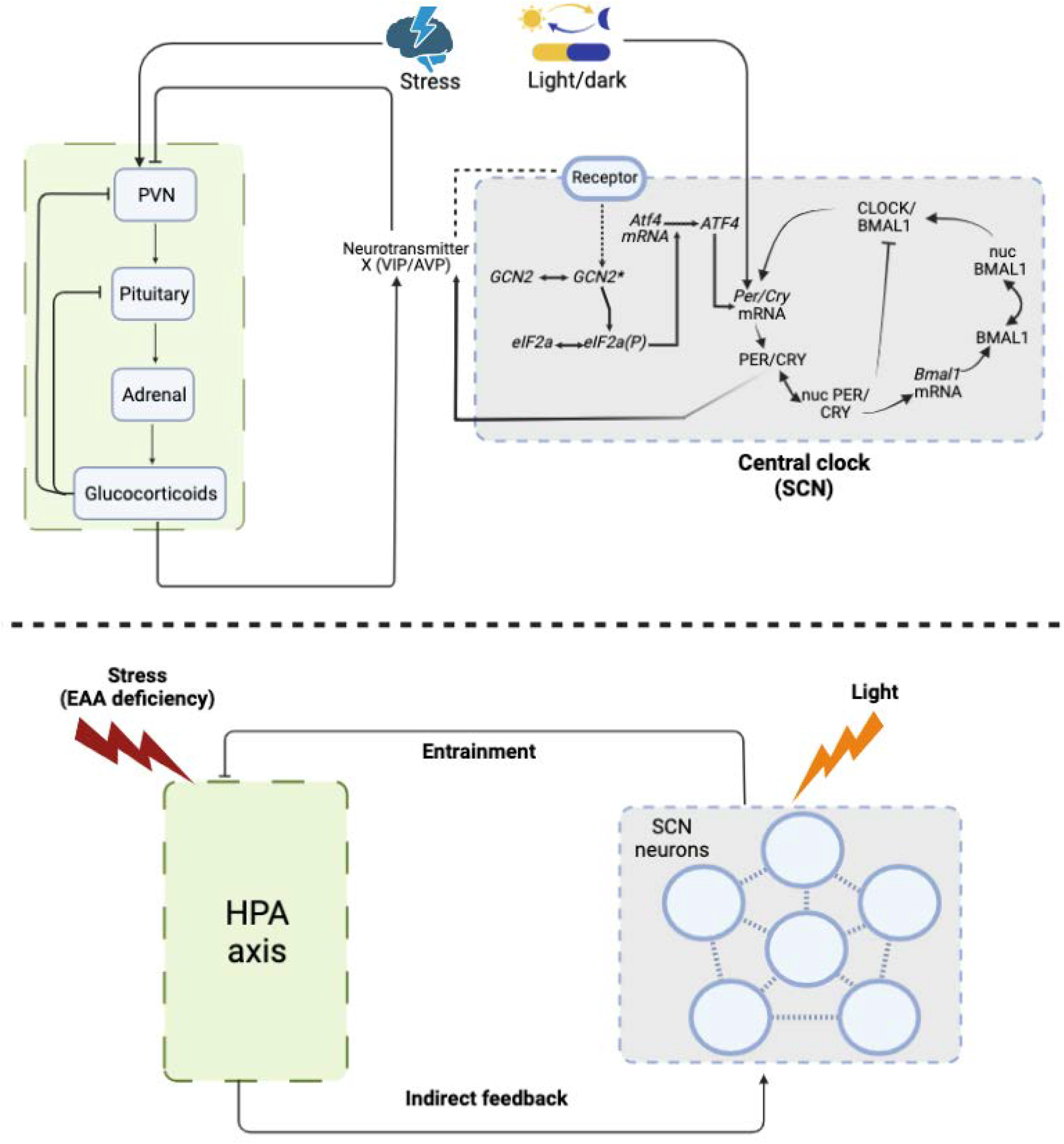
Schematic representation of the interaction between the Integrated Stress Response (ISR) sensing system and circadian rhythms. The physiological stress is represented by EAA deficiency, which is sensed by the hypothalamus. Activated by stress, the HPA axis transduces the stress information to glucocorticoids, which indirectly feedback to SCN by activating the expression of neurotransmitters. The SCN is mainly regulated by light, while the HPA axis is mainly activated by stress signals. The communication between the SCN and the HPA axis facilitates the coordination between integrated stress response and circadian physiology

The SCN functions as a network of heterogeneous neurons, each operating as an individual oscillator with a complex interplay of clock genes and proteins forming feedback loops. These cellular oscillations are mathematically detailed in our **Supplementary Materials** (Geier et al. 2005). We account for the crucial role of the eIF2α-GCN2-ATF4 pathway in ensuring robust oscillations in every individual neuron within the SCN (Pathak et al. 2019). Specifically, as one of the most evolutionarily conserved and abundant eIF2α kinases in the brain, GCN2 is required for ISR activation within the brain, with GCN2^∗^ denoting its activated state (**Equation 2**). This activation subsequently promotes eIF2a phosphorylation (**Equation 3**), facilitating the translation of transcriptional modulators such as activating transcription factor 4 (ATF4) (**Equation 4**). Through its binding to the *Per2* promoter region, ATF4 enhances the transcription of clock genes in the SCN (**Equation 5**). Consequently, the ISR sensing pathway is embedded within each SCN cell, influencing its rhythmic behavior.

In alignment with our previous research (Li and Androulakis 2022; Li and Androulakis 2023), the SCN model in the current study comprises a diverse population of neurons that release neurotransmitters (denoted as “V”). These neurotransmitters facilitate self-coupling and inter-neuronal communication within the neuronal network. The release of neurotransmitters into the extracellular medium is triggered by the activity of PER/CRY proteins, as outlined in **Equation 6**. Their role as inter-cellular coupling signals is characterized by a distance-dependent effect, meaning that neurons adjacent to the releasing neuron are more significantly influenced, as detailed in **Equations 8-9**. Specifically, the entry of coupling signals into each neuron (including the neuron itself and the adjacent neurons that affect the neuron) is designated as ‘Q’. This process is proportional to two factors: the inter-neuronal coupling strength (represented by ‘K’) and the strength of the coupling signal (‘F’), as described in **Equation 7**. The strength of these signals is determined by the average concentration of neurotransmitters, which is referred to as the local mean field (as per **Equation 8**), release by adjacent cells within a specified threshold distance (d), detailed in **Equation 9**.

Furthermore, experimental evidence suggests that GCN2 phosphorylates eIF2α in the SCN in a rhythmic pattern (Pathak et al. 2019). However, the origin of the oscillation in GCN2 activity is not fully understood. To account for this in our model, we assumed that the oscillatory behavior of GCN2 is driven by the local mean field neurotransmitter signaling between SCN neurons. The sensed neurotransmitter signal in turn activates GCN2, as shown in the first term of **Equation 1**. As a result, SCN neurons are coupled through distance-dependent coupling and sense stress signal through GCN2.

The SCN coordinates the rhythmic activity of the HPA axis, primarily through the release of AVP within the paraventricular hypothalamic nucleus (Kalsbeek et al. 2012; Kalsbeek et al. 2010). To capture the inherent oscillatory behavior of the HPA, we utilized an established HPA axis model from prior works (Mavroudis et al. 2014). This model was further refined by integrating synchronization cues originating from the SCN, as detailed in (Li and Androulakis 2021, 2022). The exact mechanism through which the SCN receives ISR signals and activates GCN2 pathways within its neurons remains unclear. However, considering that the HPA axis is the major stress axis and that glucocorticoids are believed to activate the expression of VIP in the SCN, similar to the light/dark entrainment (Larsen et al. 1994), we simulated the SCN’s indirect detection of ISR via the stress signaling of the HPA axis. In the first term of **Equation 11**, we show that CRH is activated upon exposure to physiological stress, and the effect of stress is transduced to the output of the HPA axis, i.e., glucocorticoids. CORT, in turn, activates the expression of the neurotransmitter (**Equation 6**). Since VIP activates GCN2, our model indicates that physiological stress indirectly activates GCN2 through the HPA axis. Parameter *kfb* denotes the strength of the feedback loop between CORT and VIP expression in the SCN.

In the model, light exposure to SCN neurons is depicted as a step function across a 12-hour photoperiod, with Zeitgeber time (ZT) 0-12 marking daylight and ZT 12-24 as darkness (**Equation 1**). Physiological stress on the HPA axis is modeled in relation to this light cycle; circulating stress signals alternate between double the baseline for 12 hours and the baseline level. For nocturnal species, heightened stress either aligns with their active dark phase (ZT 12-24), or with ZT 0-12 for rest-phase stress (**Equation 10.1-10.2**). Acute stress is portrayed as a fourfold increase over baseline for 3 hours, then returning to baseline (**Equation 10.3**). This model offers a conceptual approach to understanding the influence of physiological stress on the HPA axis.

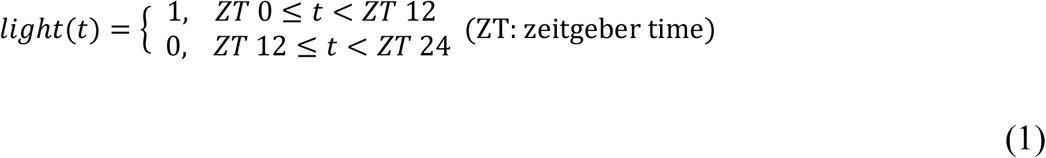

### Single cell model in the SCN

(the symbol “*i*” represents the index corresponding to a specific neuron):

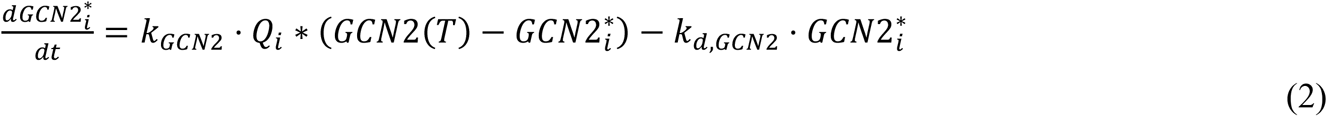

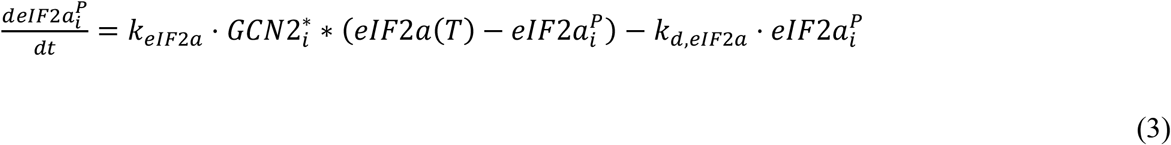

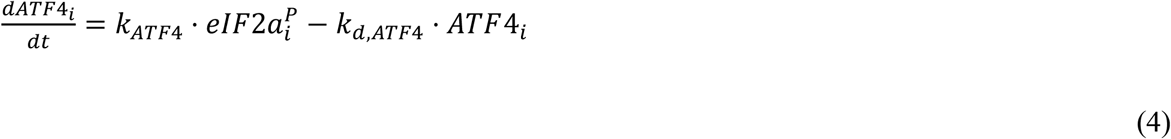

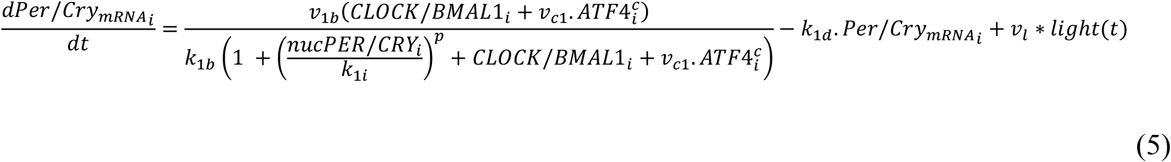

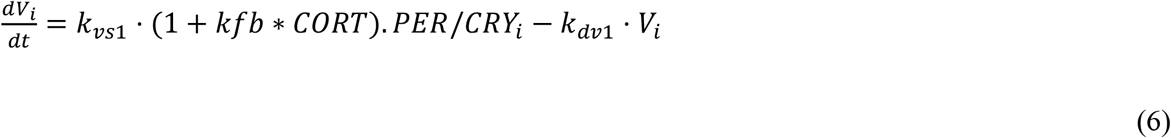

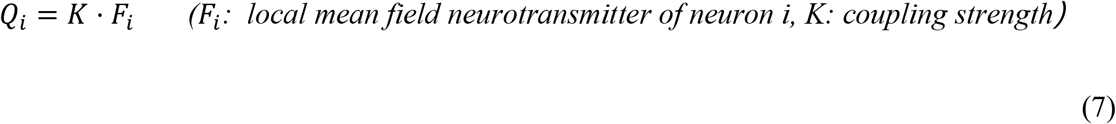

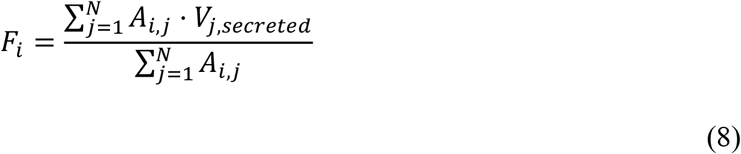

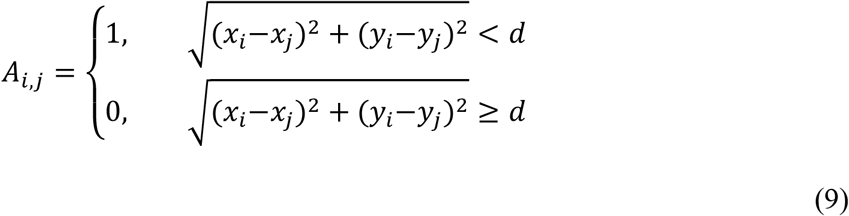

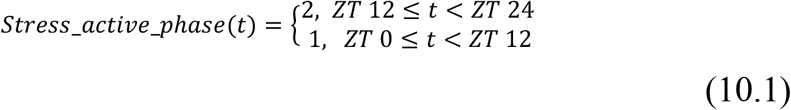

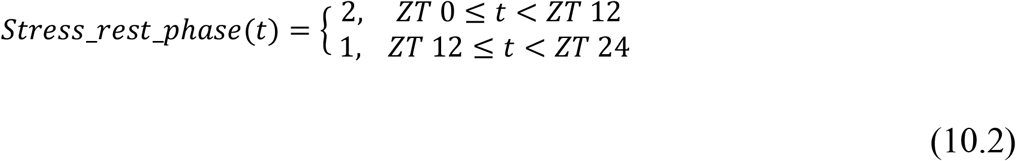

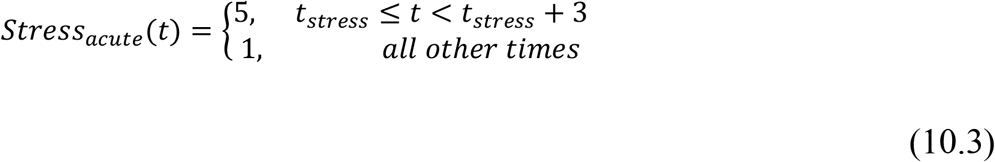

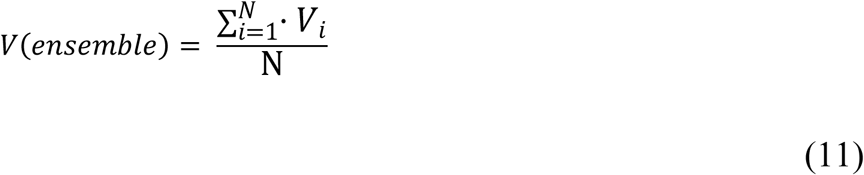

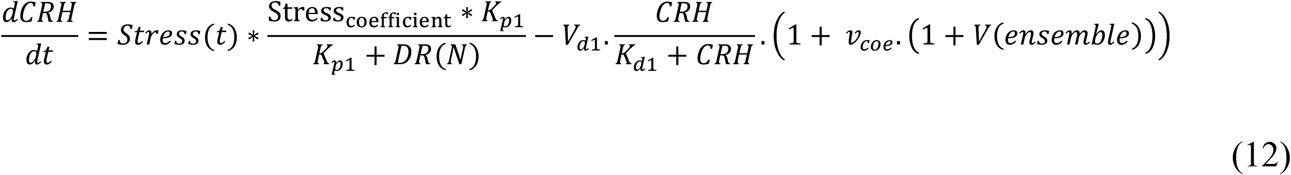

## Results

### Significance of the GCN2-eIF2α-ATF4 ISR Sensing Pathway in Sustaining Robust Oscillations within the SCN

Using our model, we performed simulations to investigate the synchronization behaviors within the SCN compartment, as illustrated in **Figure 2**. In scenarios where neurons were uncoupled (**Fig. 2a**), a lack of neurotransmitter signals from the adjacent neurons resulted in attenuated oscillations at both individual and ensemble scales. In the self-coupled scenario (**Fig. 2b**), the SCN neurons were solely influenced by the neurotransmitter signals they themselves produced. The self-secreted neurotransmitter formed an internal loop, facilitated by the ISR pathway, led to enduring oscillations. However, without mutual signals between neurons in this scenario, there was a lack of phase synchronization at the ensemble level.

**Figure 2.**
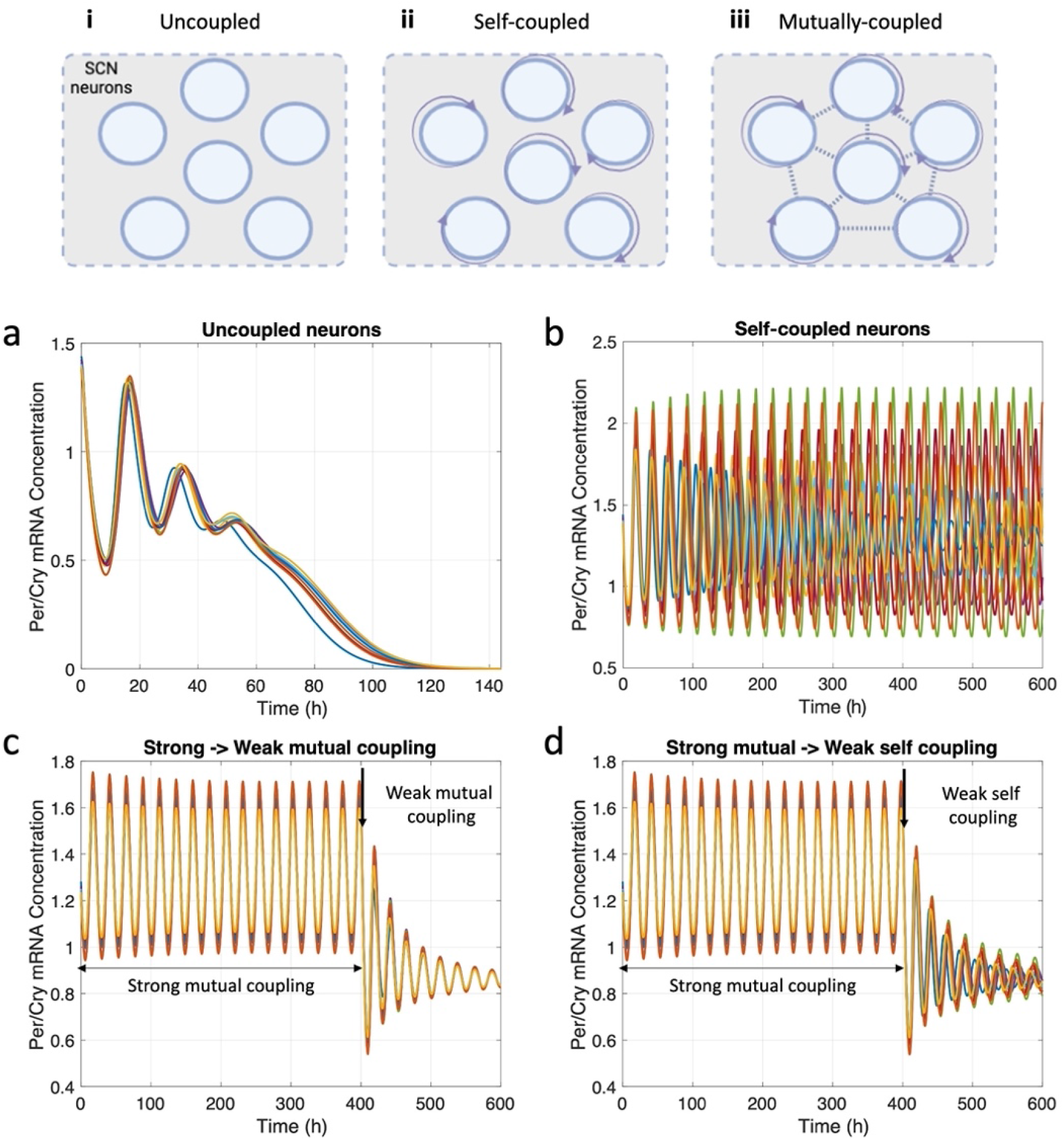
Rhythmic behavior of SCN neurons in response to coupling variations. The simulations were conducted on an ensemble of the SCN consisting of 500 cells. To visually demonstrate the findings, results from 10 of these cells are plotted. It is important to note that the selection of these particular cells for illustration does not affect the conclusions derived from the figure. The upper panel illustrates the three coupling scenarios: **i** uncoupled, where SCN neurons do not receive neurotransmitter signals from other neurons or themselves; **ii** self-coupled, where each neuron only receives its own secreted neurotransmitter signal; **iii** mutually coupled, where neurons perceive neurotransmitter signals from both themselves and other neurons. The coupling intensity is governed by the parameter K, the coupling coefficient. The bottom panel depicts the SCN Per/Cry mRNA synchronization patterns: (a) damped or absent oscillations in the uncoupled state, (b) sustained oscillations with phase desynchronization in the self-coupled state, and (c-d) coherent oscillatory dynamics among the SCN in the mutually coupled state. (c) A reduction in the coupling coefficient leads to collective damping of SCN neurons. (d) If signals from other neurons are blocked while weakening the coupling, individual neurons show semi-consistent oscillations, but their phases become asynchronous.

Transitioning to mutually coupled scenarios (**Fig. 2c-d**), neurons were interlinked, enabling them to sense neurotransmitter signals from both themselves and their neighbors. The magnitude of this mutual interaction was governed by the coupling coefficient, *K*. As *K* diminished, oscillations at the ensemble scale gradually became less robust (**Fig. 2c**). In situations where the coupling was weakened with external neuronal signals being excluded, the individual neurons showcased semi-consistent oscillations with phases diverged over time (**Fig. 2d**). These findings underscore the importance of the ISR sensing pathway in modulating SCN oscillations, especially in synchronizing the neurons and maintaining their robustness.

To delve deeper into our hypothesis emphasizing the pivotal importance of the ISR sensing pathway in the synchronization of SCN neurons, we ran simulations evaluating the amplitude and period of *Per/Cry* expression in the SCN under GCN2 knockout (KO) conditions. **Figure 3** illustrates the circadian dynamics observed under light/dark entrainment across three scenarios: wild-type (WT), GCN2 partially reserved (where GCN2 levels are reduced to half of its nominal value), and full knockout (where GCN2 levels are completely nullified). 500 SCN neurons were simulated. In **Figure 3**, the dynamics of individual neurons are depicted by grey curves, while the red curve represents their ensemble average. The circadian amplitude, as indicated on the plot, is calculated from the peak and trough of this ensemble average. In the WT setting, we noted a pronounced and stable oscillation, with the Per/Cry mRNA levels peaking during the light phase. As we increased the extent of GCN2 knockout, there was a discernible decline in the robustness of the oscillation, reflected by a diminished ensemble amplitude. Complete knockout of GCN2 led to a marked reduction in amplitude and a steeper circadian curve for Per/Cry, hinting at a compromised neural oscillator. This complete eradication of intrinsic oscillation, resulting in a very small ensemble amplitude, rendered the system acutely sensitive to variations in the light/dark cycle.

**Figure 3.**
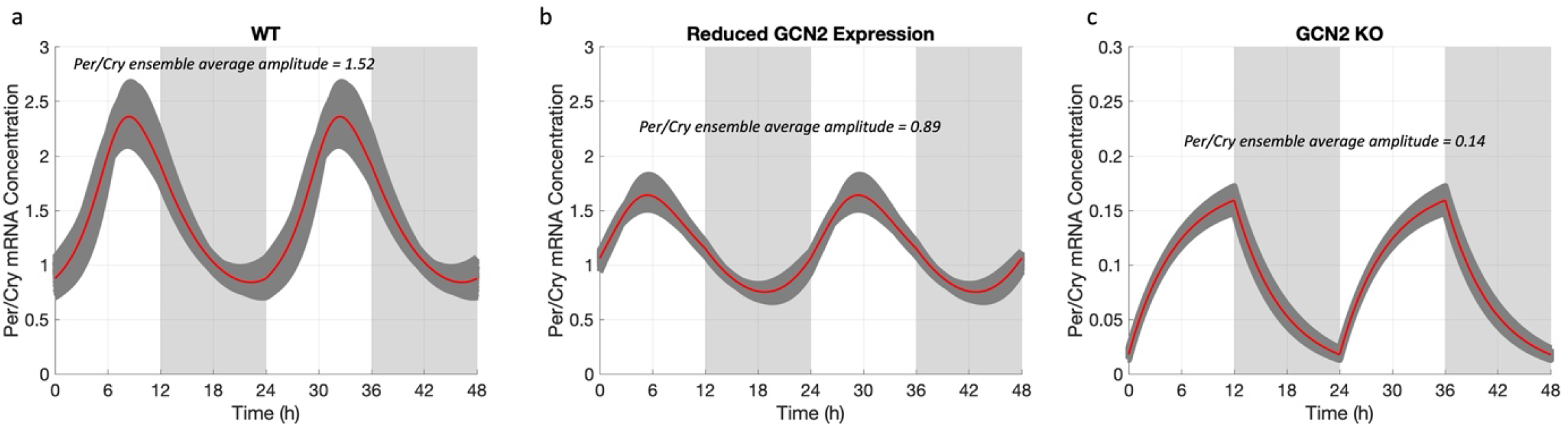
SCN neuron circadian dynamics across varied GCN2 expression conditions. Simulations were run under LD conditions, and the shadow patterns denote darkness. (a) Under LD entrainment in wild type mice, the Per/Cry displays a strong oscillation that peaks during the daytime. (b) With half the nominal GCN2 amount under LD entrainment, there’s a noticeable decrease in neural amplitude accompanied by a minor phase shift forward. (c) For SCN neurons under LD entrainment with complete GCN2 knockout (KO), a marked drop in amplitude is observed, and the Per/Cry circadian curve sharpens. This signifies a weakened neural oscillator that fully aligns with the light/dark cycle signals.

Next, we conducted simulations to investigate the period of SCN neurons and assess the effects of GCN2 knockout on circadian period. As depicted in **Figure 4a**, a decline in the total GCN2 protein level hampers the neurons’ ability to maintain oscillation, leading them to dampen swiftly. Intriguingly, a phase advance occurs at the beginning of the damping, suggesting an initial dip in the period. While this may seem to conflict with studies showing that inhibiting GCN2 and eIF2a pharmacologically extends the circadian period (Pathak et al. 2019), it’s crucial to recognize that a damped oscillator’s period evolves through its oscillation cycle. In **Figure 4b**, we plotted the SCN periods during damping across varying GCN2 levels. The findings reveal an initial shortening of the damped oscillatory period, followed by an elongation. The longer a single oscillation lasts, the fewer oscillation cycles there will be in the plot. The initial period contraction’s magnitude directly correlates with the subsequent expansion rate. For the most sustainable damped oscillation—where GCN2 is at 40% of its nominal amounts—the period initially shrinks, then rises surpassing its original length. Hence, our simulations align with and shed further light on experimental observations (Pathak et al. 2019), reinforcing the notion that the GCN2 KO system can cease its oscillation while fluidly adjusting its period. Collectively, our simulation results underscore the critical role of the ISR sensing pathway, particularly via GCN2 knockout, in modulating the SCN’s oscillatory patterns and period.

**Figure 4.**
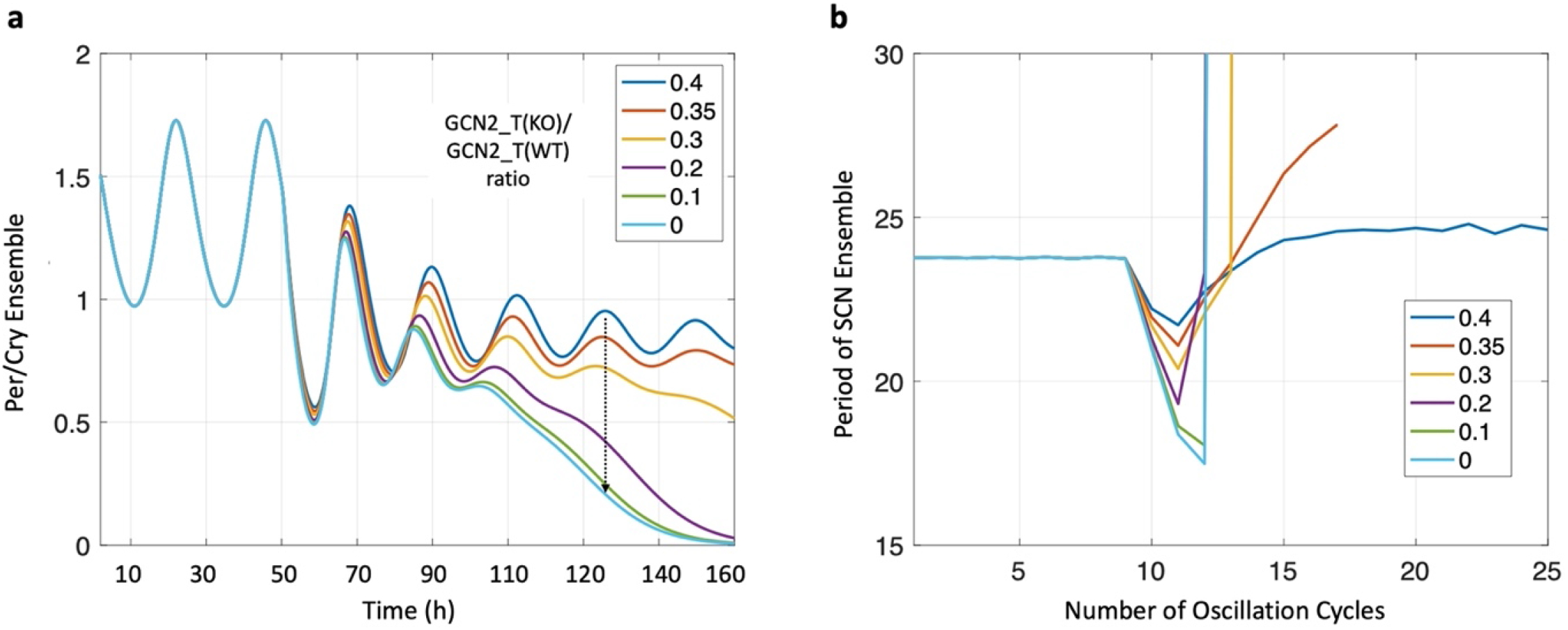
Impact of GCN2 KO on the circadian attributes of SCN neurons in constant darkness conditions. (a) A diminishing total amount of GCN2 correlates with an enhanced dampening in SCN neurons, resulting in a rapid cessation of oscillation. (b) The oscillation period of damped neurons varies over time, initially shortening, then lengthening. In scenarios where oscillation persists longer (e.g. GCN2_T(KO)/GCN2_T(WT) ratio of 0.4), the period first contracts and subsequently expands, exceeding the WT period duration. For neurons whose oscillations dampen swiftly, a notable period elongation is observed after approximately 2 days.

### Phase Response to Environmental Entrainment

Experimental data show that under standard light-dark cycles, the SCN in rodents doesn’t adjust to stress related cues such as temperature or fasting–feeding (Tahara and Shibata 2018).

But, in environments absent of light-dark cycles, like constant darkness, the SCN and the associated locomotor activity rhythm attune to temperature or feeding rhythms. Those variety of arousal stimuli can act as synchronizers for the SCN clock, referred to as “non-photic entrainment” (Buhr, Yoo, and Takahashi 2010). Since both photic and stress-induced non-photic entrainment were included in the model, we evaluated the model’s response to both entrainers by determining its phase response curve (PRC) under different entrainers. A 3h simulated light stimulus with 0.5 times intensity of the nominal light intensity and a simulated acute stress stimulus that was 5 times of the nominal intensity were introduced at different subjective time (see **Equation 10.3**), after the system had acclimated to constant darkness (DD). Acute stress was modeled as a potent trigger for CRH secretion, emulating the HPA axis’s physiological reaction to stressful events. In a DD environment, devoid of the light-dark Zeitgeber time (ZT), we earmarked the corticosterone peak as circadian time twelve (CT12). This was in alignment with experiments that set the subjective time to twelve at the onset of nocturnal species’ active phase. CT12 is hence marked by the commencement of daily activity (Hut and Beersma 2011).

Our findings, depicted in **Figure 5**, underscore a more pronounced phase response to light stimuli as compared to stress stimuli. In **Fig. 5a**, the PRC generated aligns with *type I (Daan and Aschoff 2001)*, where both phase advancements and delays can be successively initiated based on the timing of the perturbation. Intriguingly, the phase responses to light and stress display an inverse phase relationship. In **Fig. 5b**, an interesting observation is that as we increase the feedback coefficient value between the SCN and HPA, there’s a noticeable increase in the area under the curve (AUC) induced by stress stimuli. In normal physiological conditions, where glucocorticoid feedback is low, it’s inferred that both light and stress/food entrainers significantly impact peripheral rhythms (Sunderram et al. 2014) but not the SCN. This is because the area under the curve (AUC) of the stress-induced phase response curve (PRC) is minimal with a small feedback coefficient (*kfb*) between glucocorticoids and the SCN. As the feedback coefficient increases, our simulation indicates that stress stimuli may amplify the SCN’s circadian phase shift, moving it in a direction counter to that induced by light.

**Figure 5.**
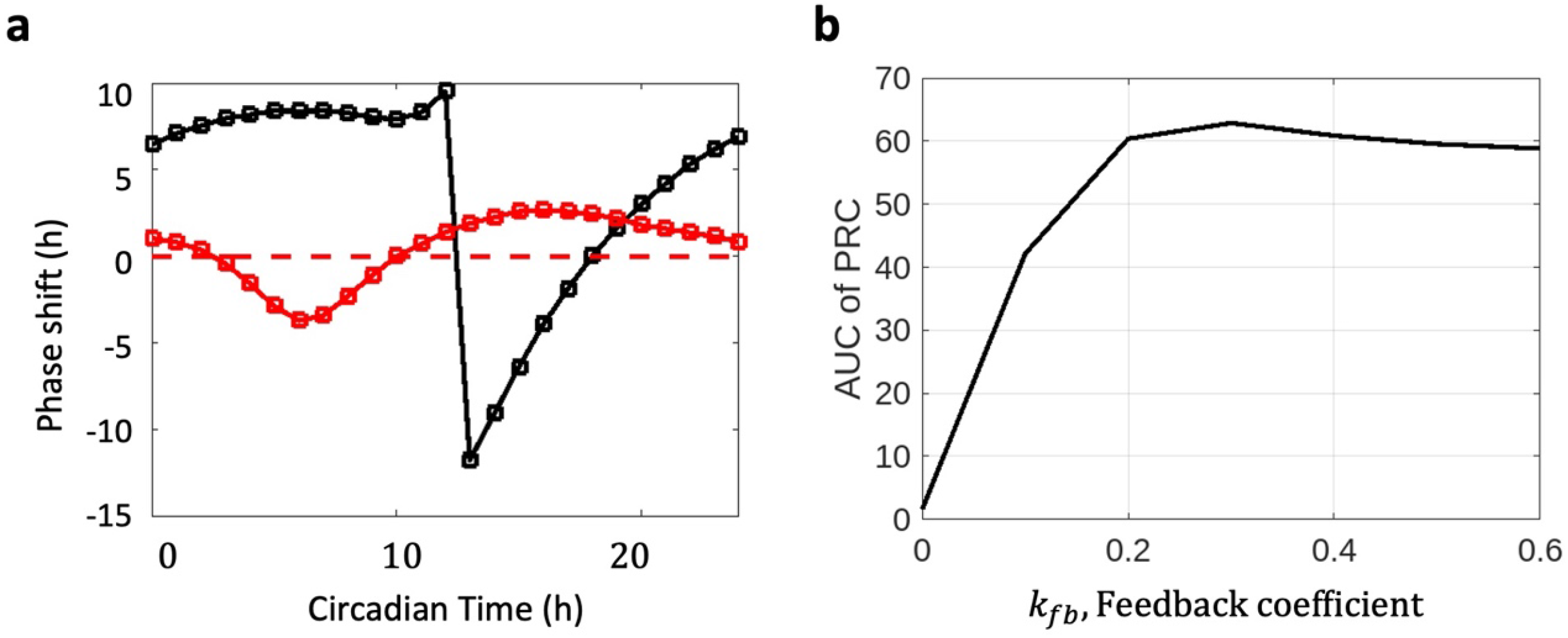
Phase Response Curve (PRC) to light and stress stimuli in constant darkness (DD) conditions. (a) PRC depicting the differential phase shifts in response to light (black) and stress (red) stimuli, showcasing an inverse phase relationship. Type I PRC indicates both phase advancements and delays based on perturbation timing. (b) Variation in the Area Under Curve (AUC) of PRC induced by stress stimuli with changes in the feedback coefficient (*k*_*fb*_) between the SCN and HPA, highlighting the increasing influence of stress stimuli. The results emphasize the contrasting roles of light and non-photic (food/stress) entrainers in regulating the SCN in different environmental conditions.

### GCN2 knockout as an intervention for jetlag

Experimental findings revealed that GCN2^-/-^ mice exhibit rapid entrainment to a shifted LD cycle, along with impaired behavioral rhythmicity and damped PER1 and PER2 rhythms in the SCN (Pathak et al. 2019). These observations led us to speculate that the accelerated jetlag transition in GCN2^-/-^ mice might be attributed to the reduced robustness of their oscillation. To investigate this hypothesis, we employed our model to simulate the jetlag behavior in both WT and GCN2 KO scenarios. **Figure 6** illustrates the phase transition process in a double-plotted actogram, showing the peaking phase for the WT system and GCN2^-/-^ system. The hosts were initially entrained to a 12h/12h light/dark cycle for 20 days, and on the 21st day, the LD cycle was advanced by 6 hours. Remarkably, the GCN2 KO mice exhibited a rapid phase shift, with their phase completely shifting to the new schedule within 1 day, whereas the WT system experienced a slower transition, taking approximately 6 days to fully adapt. Our simulated jetlag readaptation results align well with the experimental data, which reported a 1-day shift for GCN2 KO mice and a 7-day shift for WT mice to adapt to the 6-hour phase advance jetlag schedule.

**Figure 6.**
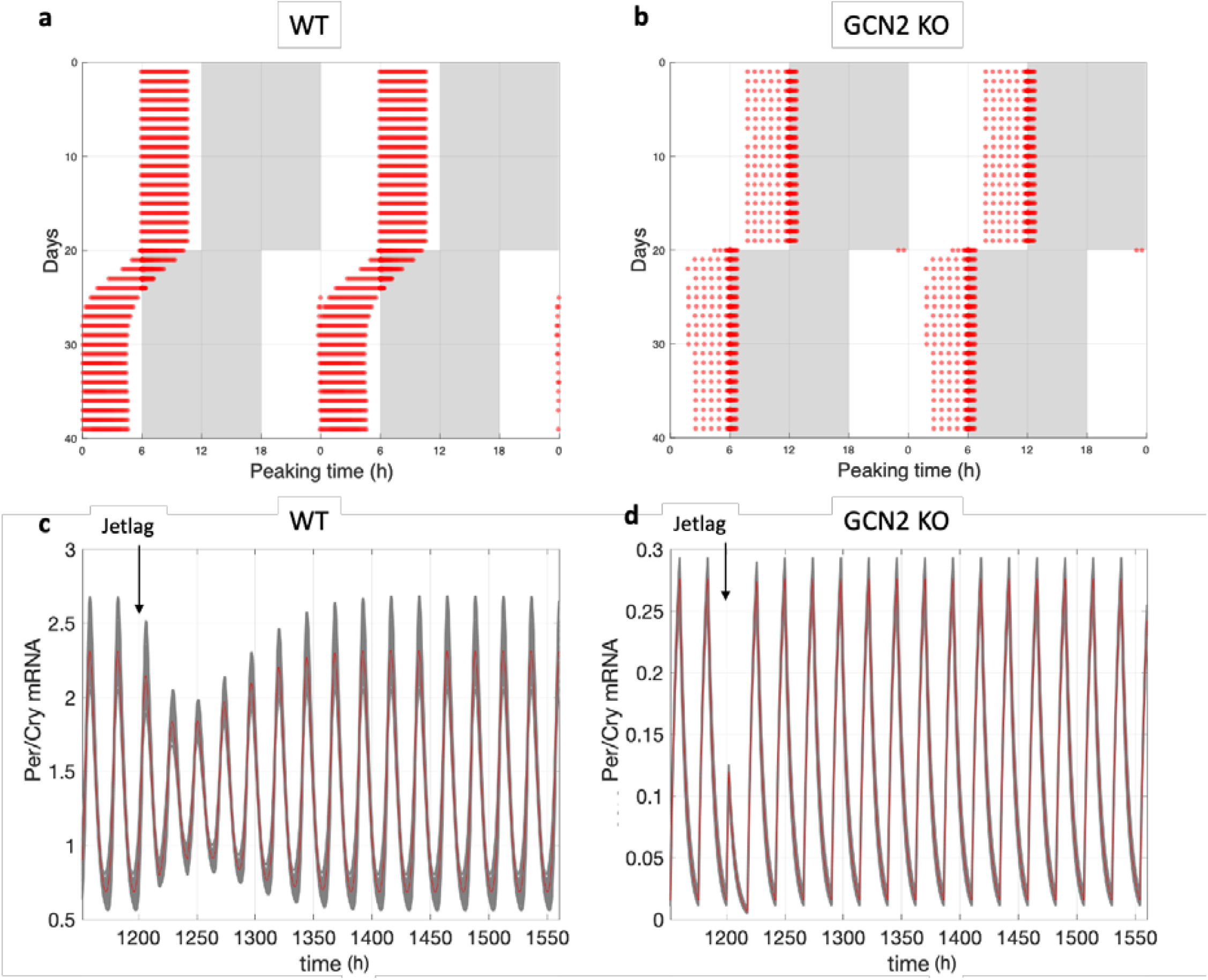
Double-plotted actograms of peaking phase from (a) WT system and (b) GCN2^-/-^ system during jetlag. The x-axis indicates the zeitgeber time of the day, and the y-axis indicates the number of days. The hosts were entrained to a 12h/12h light/dark cycle for 20 days, and on the 21st day, the LD cycle was advanced by 6 hours. (c) and (d) show the circadian dynamics of Per/Cry mRNA for WT and GCN2 KO systems under jetlag, respectively. Jetlag was imposed at 1200 h. A rapid phase shift was observed in the GCN2 KO case, while the WT system experienced a slower transition.

**Figure 6.**
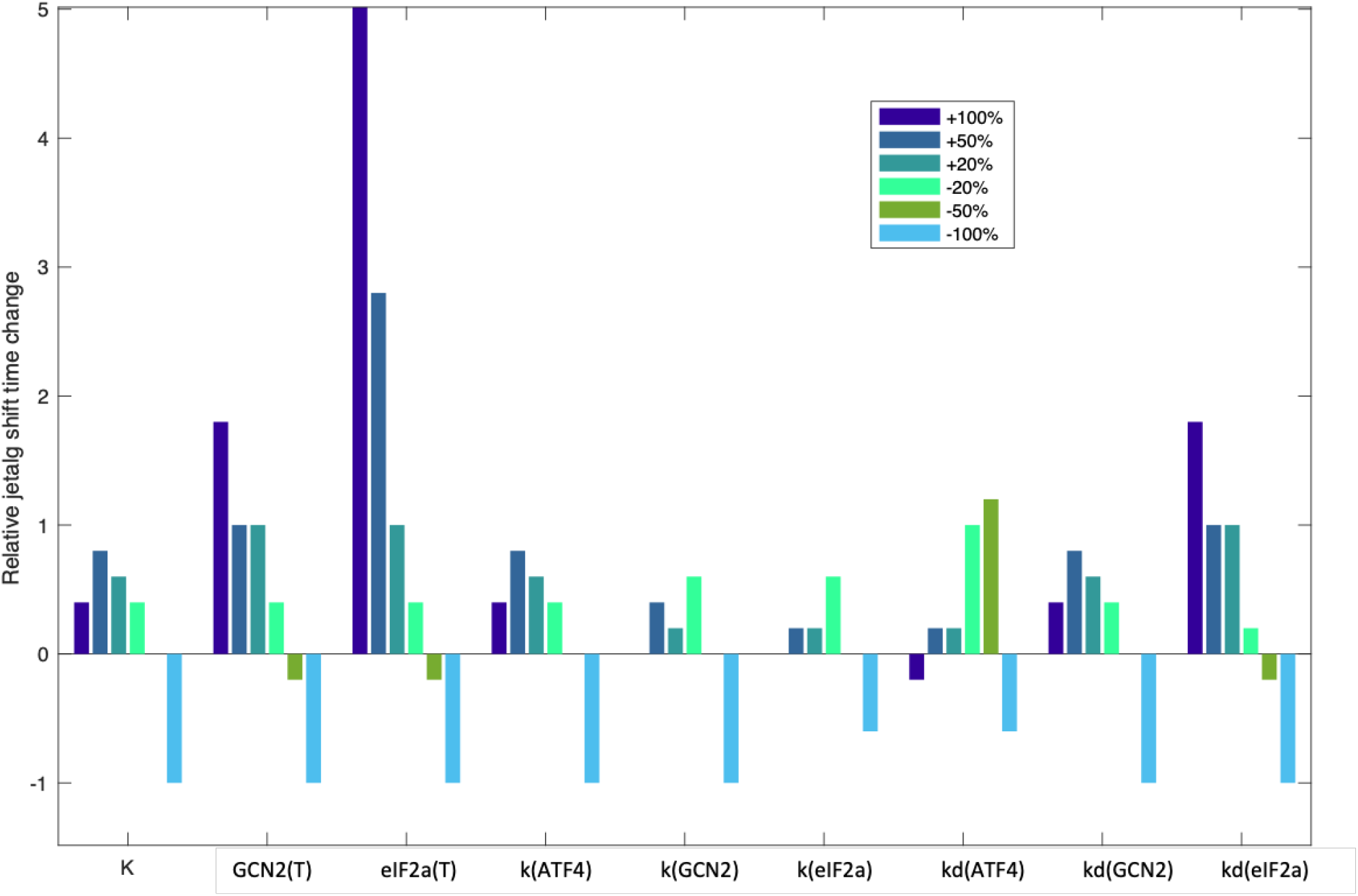
Parameter sensitivity analysis of the parameters involved in the ISR sensing pathway. The jetlag transition time was tested with different levels of varying parameters. The x-axis shows the varying parameters, while the y-axis shows the relative change in jetlag transition time compared to the nominal parameter set. The varying percentage ranges from -100% to +100% compared to the nominal value. The results demonstrate the critical role of the ISR sensing pathway in determining the jetlag re-entrainment rate. Specifically, the total amount of GCN2 and eIF2α has a significant impact on the transition rate of the system.

To quantitatively investigate the influence of different effectors in the ISR sensing pathway on the system’s jetlag recovery, we conducted a parameter sensitivity analysis on factors including neurotransmitter coupling strength and the total cellular concentrations of GCN2 and eIF2a. Additionally, we examined the sensitivity of the synthesis and degradation rates of eIF2*a*, GCN2, and ATF4, all of which are integrated to the newly added ISR pathway. We calculated the jetlag transition time with varying levels of these parameters, ranging from -100% to +100% compared to their nominal values. By comparing the transition time after parameter variation to the transition time under the nominal parameter set, we determined the most influential factors affecting the jetlag transition rate.

In **Figure 7**, our sensitivity analysis revealed that the total amount of GCN2 and eIF2α are the most significant factors in modulating the jetlag transition time. Increasing these parameters resulted in a longer transition time during jetlag, while decreasing them significantly shortened the time required for the system to adapt. Additionally, other properties, such as decreasing the coupling coefficient (*K*), deactivating ATF4 or GCN2 (decreasing *k*(*ATF*4) and *k*(*GCN*2)), and reducing the degradation rates of GCN2 and eIF2α (*kd*(*GCN*2) and *kd*(*eIF*2*α*)), all disrupted the rhythm and accelerated the jetlag recovery time.

**Figure 7.**
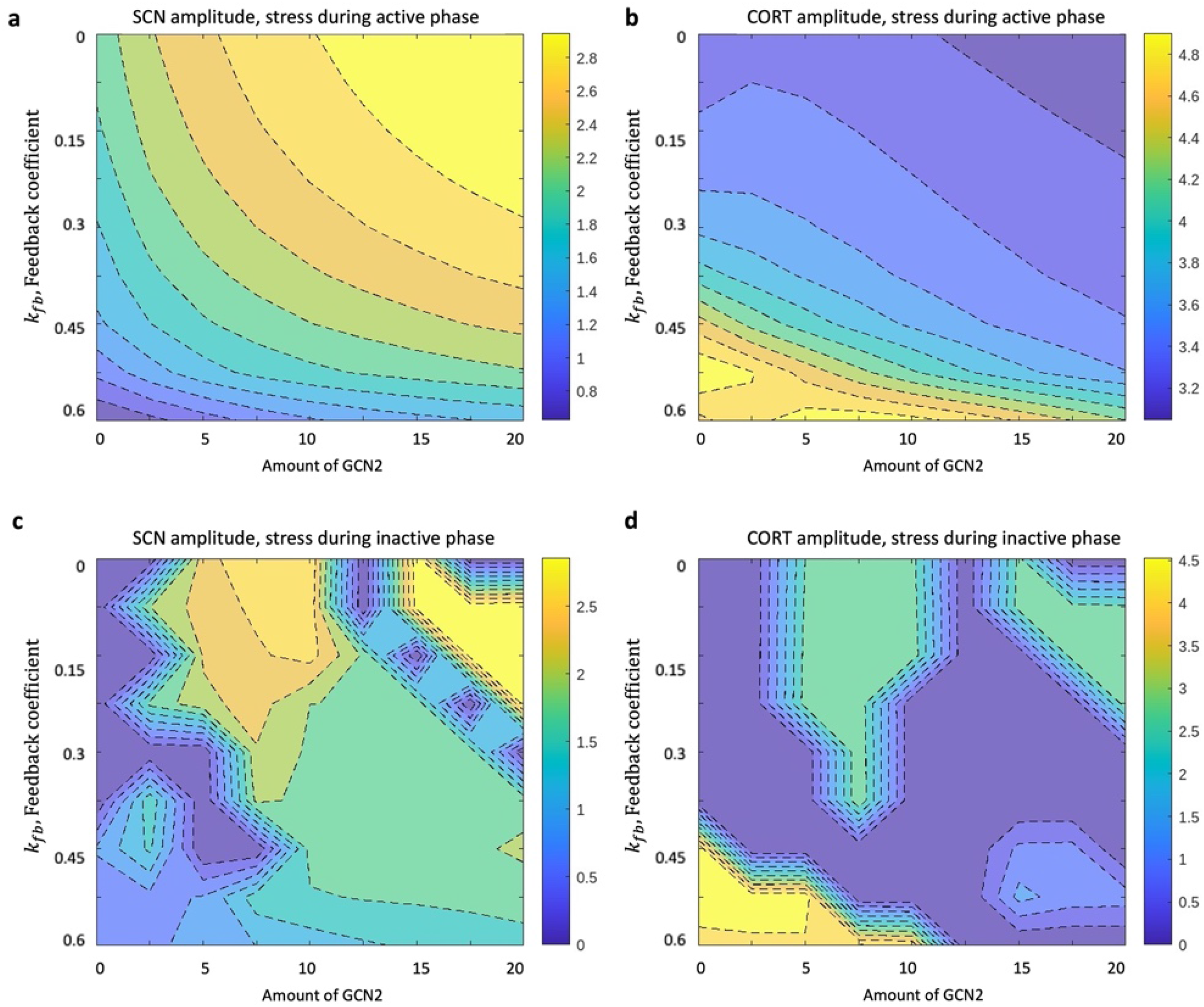
Circadian amplitude variations of the SCN and glucocorticoids under different GCN2 levels and glucocorticoid feedback strengths. (a-b) Stress alignment with the host’s active phase reveals diverse entrainment patterns in both the SCN and the HPA axis, contingent on GCN2 amounts and feedback strength. An increase in GCN2 results in elevated SCN amplitude but decreased HPA axis amplitude, suggesting that GCN2 activation can potentially assume some of the HPA axis’s responsibilities in stress signal response. Concurrently, as feedback strength surges, the SCN amplitude similarly rises, whereas the HPA axis amplitude diminishes, illustrating the synchronized response of the SCN and HPA axis to stress.(c-d) When stress transpires during the host’s non-active phase, the amplitude of both the SCN and HPA axis reduces, leading to irregular entrainment patterns. This indicates a compromised efficiency of the host in managing stress during the inactive phase compared to the active phase. Optimal homeostasis is achieved when stress cues are presented during the active phase. The model further intimates that amplified SCN oscillations are mainly discernible in zones with a minor feedback coefficient, suggesting an upper limit to the coefficient size for ideal SCN synchronization with the light-dark cycle.

In summary, our results highlight the crucial role of the ISR sensing pathway in determining the rate at which the system re-entrains after jetlag. Specifically, the total amount of GCN2 and eIF2α exert a significant impact on the transition rate of the system.

### The interplay between circadian physiology and integrated stress response

Aligned with experimental results, our model thus far strongly supports the notion that the ISR sensing pathway is instrumental in maintaining the oscillatory behavior of the SCN by modulating the robustness of its intrinsic neural oscillators. Building upon this, we delved deeper into our hypothesis regarding the stress-sensing function of the ISR sensing pathway in the SCN, in conjunction with the HPA axis, through the indirect feedback mechanism involving CORT regulation of VIP expression. To thoroughly evaluate the effects of GCN2 levels and stress profiles on the circadian rhythms of the SCN and HPA axis cortisol output, our analysis covered circulating physiological stress scenarios. These scenarios encompass stress events that occur in a recurring manner and trigger the physiological response of the HPA axis. We examined conditions where physiological stress profiles are synchronized with the active phase of nocturnal organisms and conditions where they are out of sync with this active phase.

The circadian amplitudes of both the SCN and glucocorticoids were calculated under varying levels of GCN2 and glucocorticoid feedback strength. In **Figure 7a-b**, we demonstrate that when stress aligns with the host’s activity phase, both the SCN and the HPA axis exhibit a wide range of entrainment patterns depending on the combination of GCN2 total amount and feedback strength, as reflected by the oscillatory amplitude of Per/Cry in the SCN and glucocorticoids. While a sufficient amount of GCN2 is essential for achieving robust oscillation within the SCN, this surge might diminish the amplitude of HPA oscillation. As GCN2 levels rise, the SCN amplitude elevates, whereas the HPA axis amplitude declines. This suggests that GCN2 activation can partially take over the HPA axis’s role in responding to stress signals. Similarly, as the feedback strength increases, the SCN amplitude increases while the HPA axis amplitude decreases, suggesting that the coupling between the SCN and the HPA axis harmonizes the response of both compartments to stress.

In **Figures 7c-d**, we observe that if stress occurs during the rest phase, the amplitude of both the SCN and the HPA axis decreases, and the entrainment pattern becomes irregular. This suggests that the host’s ability to cope with stress during the inactive phase is not as effective as during the active phase. The system achieves optimal homeostasis when stress signals occur during the active phase. Furthermore, given that the amplified SCN oscillation predominantly appears in regions with a smaller feedback coefficient, the model suggests that this coefficient should not be excessively large. A diminished SCN responsiveness to the CORT signal may aid in synchronizing the SCN with the light-dark cycle.

Given the prevalent individual variability inherent to physiological systems, our model is intended to serve as an overarching representation, encapsulating a composite of diverse individuals. It stands to reason that the model’s key parameter variations can capture individual differences and elucidate correlations with distinct properties (Sterling 2012). Building on the prior insight that the GCN2, which signifies the ISR sensing pathway, collaboratively functions with the HPA in stress response, we hypothesized a potential association between an individual’s stress resilience and its GCN2 expression levels. Moreover, we were keen to discern if this relationship is influenced by the HPA-to-SCN feedback strength.

To probe this, we sampled three key parameters: the total GCN2 concentration in the SCN cells, the stress coefficient which reflects the intensity of an individual’s CRH response to stress (as detailed in Equation 12), and the feedback coefficient between glucocorticoids and the SCN, which represents the SCN’s sensitivity to glucocorticoid signaling. A virtual population was constructed using Sobol sampling, ensuring that the circadian profiles of simulated CORT remained within a ±2-hour bracket of the standard CORT peaking phase (ZT 12). This sampling strategy facilitated the identification of a parameter subspace where the CORT rhythms are in homeostasis. Upon generating this population, we gauged their glucocorticoid amplitude, denoting the resultant value through dot coloration.

Figure 8. reveals a broad acceptable distribution range for both the stress coefficient and GCN2, in contrast to a more confined range for the feedback coefficient. This suggests a vast variability in individuals’ stress responsiveness, while the SCN’s reception of signals from the HPA axis remains more constrained. Notably, a positive correlation between GCN2 expression and the stress coefficient was evident. This suggests a synchronicity in stress detection between the SCN and HPA; heightened sensitivity to stress in the HPA corresponds with increased ISR signal sensitivity in the SCN. Furthermore, as both GCN2 concentration and stress coefficient escalate, glucocorticoid levels also rise, hinting that a surge in an individual’s stress sensitivity amplifies its circadian robustness. While the feedback coefficient did not display marked correlations with other stress traits, there’s a notable clustering of individuals at lower feedback coefficient values, aligning with literature indicating the SCN’s relative insensitivity to downstream HPA signals.

**Figure 8.**
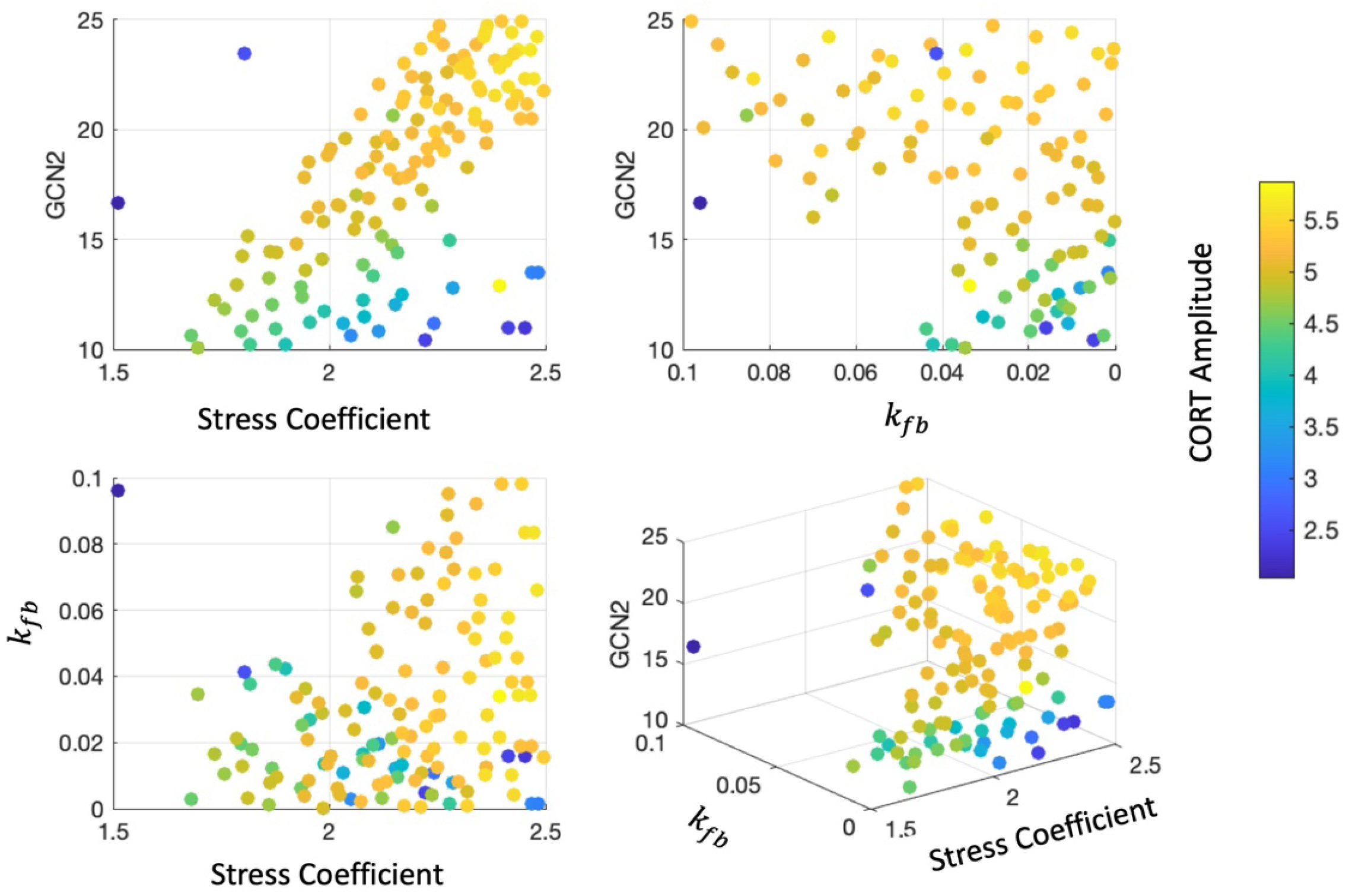
Scatter plots illustrating the distribution and relationship between GCN2 expression, stress coefficient, and feedback coefficient. 300 virtual individuals were simulated. The varied color intensity of the dots represents glucocorticoid amplitude, highlighting the correlation between stress sensitivity and circadian robustness. The charts demonstrate individual variability in stress responsiveness and the constraints in SCN’s reception of HPA axis signals.

## Discussion

The ISR sensing pathway, particularly the circadian phosphorylation of eIF2α by GCN2, has been shown in experimental studies to be crucial for sustaining oscillations in the SCN. It has been observed that rhythmic PER protein oscillations in the SCN rely on this phosphorylation process (Pathak et al. 2019). The phosphorylation of eIF2α by GCN2 in the SCN promotes *Per2* transcription by controlling the translation of *Atf4*. Studies conducted on GCN2^-/-^ mice have revealed reduced levels of both *Per1* and *Per2* in the SCN, under both constant darkness and light-dark conditions. These results suggest that GCN2 is instrumental in modulating the circadian rhythms of clock genes and proteins within the SCN and should be recognized as an essential factor ensuring oscillation when simulating SCN neurons.

On the other hand, the coupling between neurons within the SCN has been demonstrated to be essential for maintaining oscillatory behavior. Experimental and theoretical investigations have shown that synchronization factors among SCN neurons not only coordinate cellular activity but also play a critical role in sustaining intrinsic rhythmicity. Disruption of intercellular signaling leads to a loss of sustained rhythmicity in most neurons. Bernard et al. proposed a model suggesting that periodic synchronization signals are necessary for maintaining rhythmicity in the majority of SCN neurons (Bernard et al. 2007). These findings lead us to hypothesize that GCN2 is involved in the coupling and sustaining of SCN neurons, along with their intercellular synchronization.

While direct evidence of GCN2’s role in coordinating coupling between SCN neurons is lacking, similar metabolic kinases have demonstrated widespread interaction with SCN coupling. For example, in mice, the mammalian target of rapamycin complex 1 (mTORC1) has been shown to phosphorylate the translation repressor eukaryotic translation initiation factor 4E (eIF4E)-binding protein 1, regulating circadian clock entrainment and clock cell synchrony by facilitating mRNA translation of Vip (vasoactive intestinal peptide) in the SCN (Cao et al. 2013). Given the shared regulatory pathways and functions between GCN2 and mTORC1, particularly concerning amino acids (Averous et al. 2016; Ye et al. 2015; Misra et al. 2021), it is reasonable to presume that GCN2 plays a role in coupling SCN neurons. To account for the significance of eIF2α phosphorylation and subsequent transcriptional regulation in initiating the ISR, the single-cell model of the SCN incorporates key components of the ISR sensing pathway. During ISR activation, GCN2, the eIF2α kinase, becomes activated and crucially phosphorylates eIF2α. This phosphorylation prompts the translation of transcriptional modulator ATF4, thereby enhancing the transcription of Per2.

By incorporating a mathematical model that simulates the presumed mechanisms of the ISR sensing pathway in the SCN, our study successfully reproduces experimentally observed phenomena, including damped oscillations and an extended period of the SCN rhythm in the absence of GCN2. These simulation results qualitatively validate the importance of the eIF2α-GCN2-ATF4 pathway in sustaining robust oscillations within the SCN. Furthermore, our findings offer valuable insights into the intricate coupling between the ISR sensing pathway and the central circadian clock.

Our jetlag results indicate that increased levels of GCN2 and eIF2 enhance the robustness of circadian clocks, making them less susceptible to re-entrainment by external cues, akin to the prolonged re-entrainment observed in GCN2 knockout (KO) mice. Biologically, heightened GCN2-eIF2 signaling correlates with reduced protein synthesis rates (Pettit et al. 2017). In the brain, we speculated that this reduction in protein synthesis could decelerate the phase adjustment process during jetlag transitions. Conversely, the absence of GCN2 removes the usual suppression of protein synthesis in response to stress, potentially hastening the re-entrainment process. Our simulation, which targeted the ISR sensing pathway, suggests that pharmacological inhibitors aimed at GCN2 and eIF2α could serve as a potential strategy for alleviating the effects of jetlag. Nevertheless, it is essential to acknowledge that inhibiting these factors may compromise the resilience of the intrinsic oscillator. Moreover, previous studies have implicated GCN2 in governing various neurophysiological processes, such as synaptic plasticity, learning and memory, and feeding, further supporting our hypothesis that GCN2 plays a vital role in coupling SCN neurons. Disruption of GCN2 may lead to reduced robustness while increasing adaptability to external disturbances (Maurin et al. 2005; Costa-Mattioli et al. 2005).

Furthermore, the study investigated the role of the hypothalamic-pituitary-adrenal (HPA) axis in coupling the ISR signal and the central circadian clock. The HPA axis, as the major stress axis, transduced stress signals to glucocorticoids, which indirectly activated the expression of neurotransmitters in SCN neurons. This indirect activation of GCN2 through the HPA axis provided a potential mechanism for the SCN to sense stress signals. The feedback loop between glucocorticoids and the SCN was proposed to facilitate the coordination between stress response and circadian physiology.

The presented model in this study serves as a simplified representation, and further research is required to validate its predictions and explore additional mechanisms. To gain a more comprehensive understanding of the intricate dynamics, future investigations should incorporate more detailed information about the metabolic and stress signaling pathways in both the central and peripheral circadian compartments. Nevertheless, this study stands as the first theoretical work to investigate the complex interaction between ISR sensing and central circadian rhythm regulation, encompassing the SCN and the HPA axis. These findings carry implications for the development of dietary or pharmacological interventions aimed at facilitating recovery from stressful events, such as jetlag. Moreover, they provide promising prospects for potential therapeutic interventions that target circadian rhythm disruption and various stress-related disorders.

## Supplementary Materials

### 1. Materials and Methods

#### 1.1. Model Development

In this study, we have refined our model to incorporate the pivotal role of the GCN2-eIF2α-ATF4 pathway (GCN2, general control nonderepressible 2; eIF2α, eukaryotic translation initiation factor 2α; ATF4, activating transcription factor 4) within the integrated stress response (ISR) as an internal mechanism in the suprachiasmatic nucleus (SCN). This pathway is essential for the resilience of the central mammalian clock located in the SCN. Our model also highlights the interconnected reliance of this master clock’s response to metabolic stress signals on feedback from the hypothalamic-pituitary-adrenal (HPA) axis. This relationship is framed within the SCN topology-HPA axis model established in our previous work [1, 2]. Distinctively, the model differently represents the SCN as a heterogeneous collection of GCN2-eIF2α-ATF4 pathway-mediated, damped neuronal oscillators and further incorporates the hypothetic, indirect effect of glucocorticoids (CORT), which is output from the HPA axis, on upregulating the neurotransmitter expression within SCN neurons (**Figure 1**).

##### 1.1.1. The ISR pathway (GCN2-eIF2α-ATF4 signaling cascade)-mediated intra-neuronal and inter-neuronal coupling in the SCN

Based on the mechanism elucidated by Pathak and Cao et al. [3], where the ISR pathway modulates the circadian characteristics of SCN clock by regulating the transcription of clock gene (*Per2*), we integrated the GCN2-eIF2α-ATF4 pathway with the autoregulatory clock dynamics into each single neuron in the SCN. Upon the GCN2 is activated (*GCN*2^∗^) (**Equation 1**), eIF2α is phosphorylated (*eIF*2*a*^*P*^) (**Equation 2**) to initiate the ISR and promote the translation of transcriptional modulators such as ATF4 (**Equation 3**), which then enhances the transcription of *Per/Cry* mRNA (**Equation 9**) by binding to the *Per2* promoter region and modulate the clock genes and proteins dynamics. The deactivation/ dephosphorylation or selective degradation of these proteins are described using negative multipliers of protein concentrations and corresponding rate constants.

*A single cell (cell i) in the SCN:*

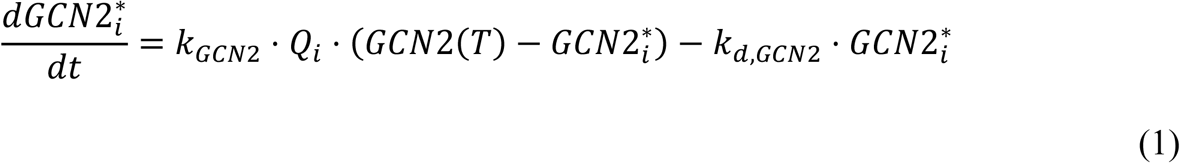

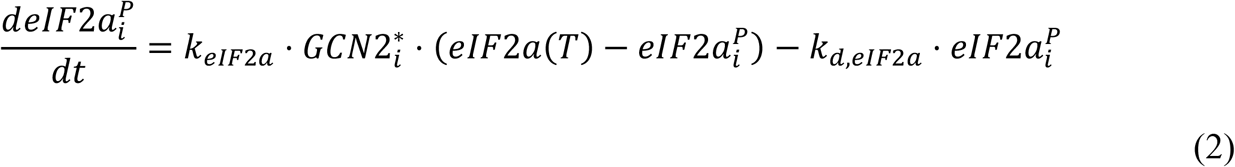

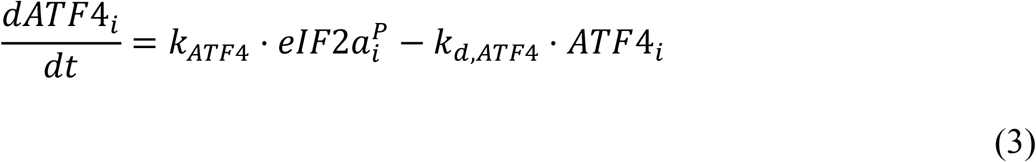

Similar to our prior works [1, 2], the SCN in this study is represented as a single compartment that consists of a population of neurons with heterogeneity, where the component SCN neurons release neurotransmitters (*V*) (**Equation 4**) for both self-coupling at the cellular level and inter-neuronal communication at the tissue level. The secretion of neurotransmitters to the extracellular medium are assumed to be induced upon the activity of PER/CRY proteins (**Equation 4**) and their functions as inter-cellular coupling signals are distance-dependent, implying that the adjacent neurons of the neurotransmitter-releasing neuron are more affected (**Equation 5-7**). Specifically, the entry coupling signals to each cell (both the same neurons and affected/ coupled neighbors) (*Q*) is proportional to the intra-SCN inter-neuronal coupling strength (*K*) and the strength of coupling signal (*F*) (**Equation 5**), which is calculated by the average concentration (local mean field) of neurotransmitters (**Equation 6**) released by all cells within the threshold distance (*d*) (**Equation 7**).

*Intra-cellular and inter-cellular coupling mechanisms of the SCN:*

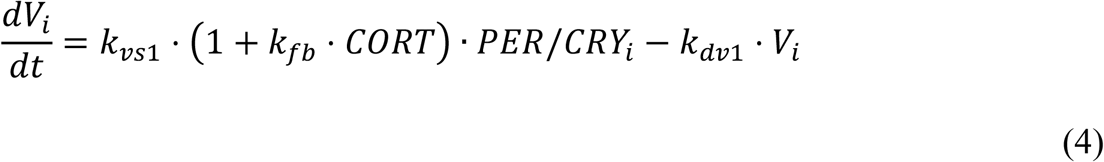

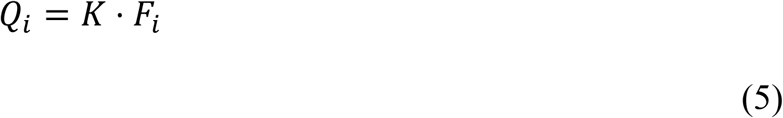

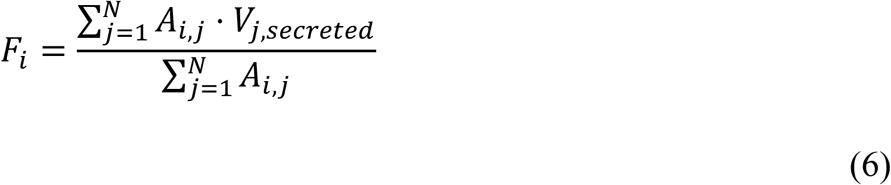

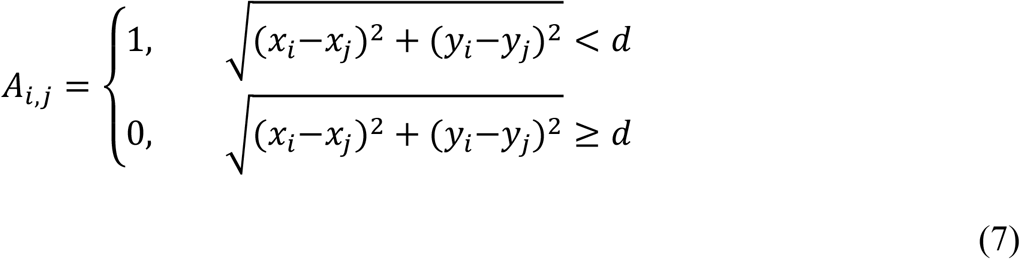

Furthermore, experimental evidence suggests that in the SCN, activated GCN2 phosphorylates eIF2α in a rhythmic pattern [3], the driving force of the oscillation in GCN2 activity, however, is not fully understood. Given the significances of neurotransmitters involved coupling mechanism and of GCN2-eIF2α-ATF4 pathway to the SCN clock’s robustness/ plasticity, we thus hypothesized that the GCN2-eIF2α-ATF4 signaling cascade, which activates the transcription of *Per/Cry*, and accordingly the *GCN*2^∗^rhythm in each cell are triggered by the entry neurotransmitter effects (the first term in **Equation 1**).

##### 1.1.2. The clock gene dynamics in the SCN neurons

The intrinsic dynamics of the clock genes and proteins network is modeled using the same gene regulatory network [4, 5] as our previous works [1, 2, 6, 7], which consists of interacted positive and negative transcriptional translational feedback loops. The positive branch is constituted by a sequence of indirect activation of *Bmal1* transcription by nuclear PER/CRY protein (**Equation 12**), translation of *Bmal1* mRNA (**Equation 13**), nuclear translocation of cytoplasmic BMAL1 protein (**Equation 14**), and heterodimerization to CLOCK/BMAL1 complex (**Equation 15**). In contrast, the inhibition of CLOCK/BMAL1-induced *Per/Cry* transcription by the nuclear PER/CRY (**Equation 9**) upon the translation to cytoplasmic PER/CRY protein (**Equation 10**) and subsequent translocation to the nucleus (**Equation 11**) forms the negative branch.

The light entraining effect on the SCN oscillators is retained as additive terms that describe the independent photic-induced *Per/Cry* transcription from the CLOCK/BMAL1-activated transcription [8] (**Equation 9**), with the 12L/12D light/dark cycle is modeled by a step function in the *in silico* experiments (**Equation 8**).

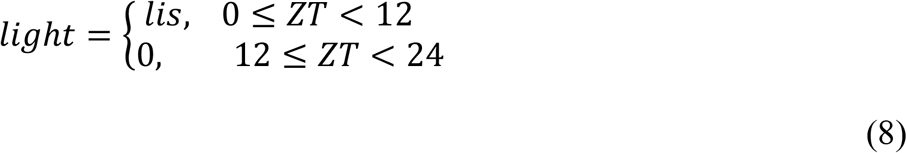

*A single cell (cell i) in the SCN:*

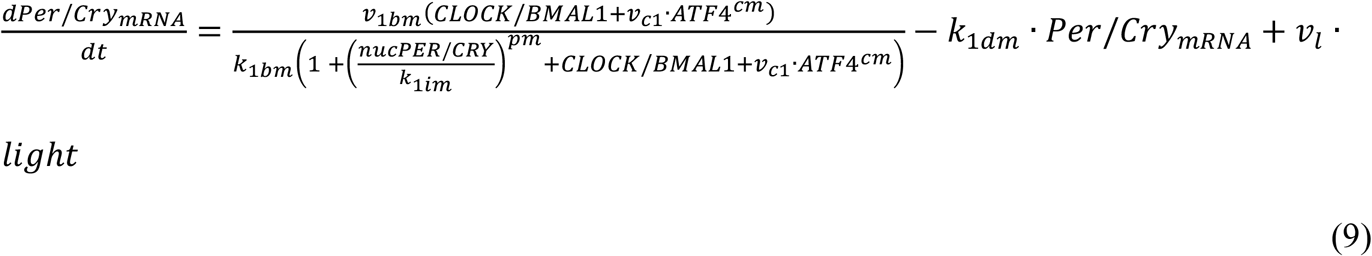

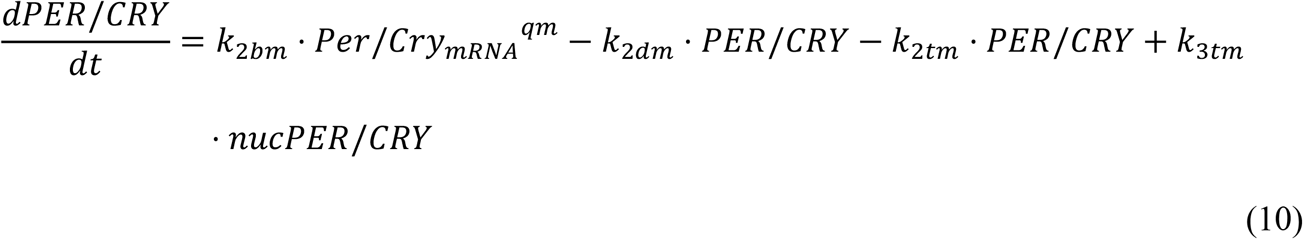

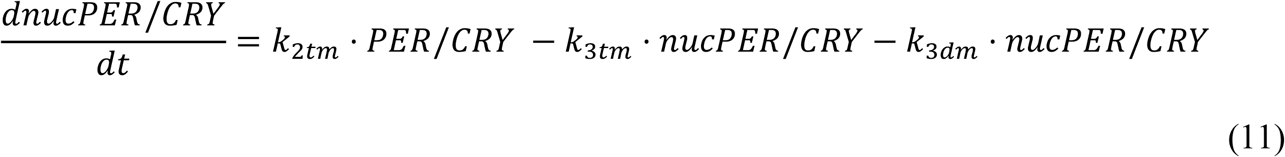

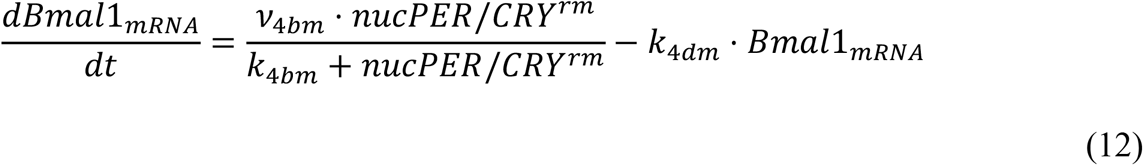

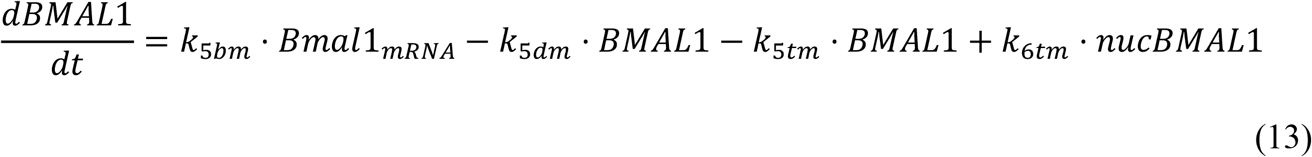

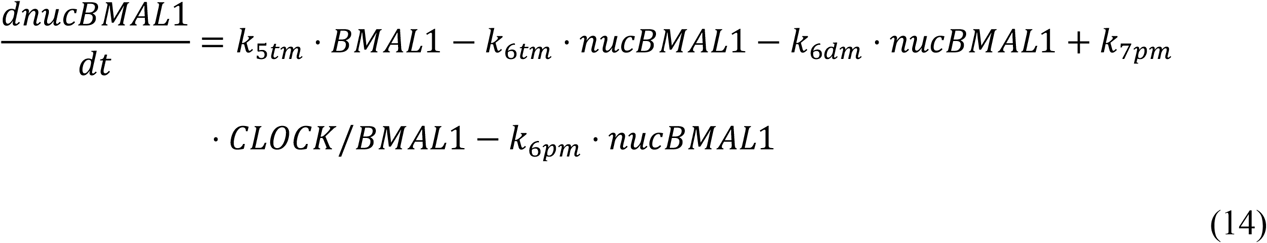

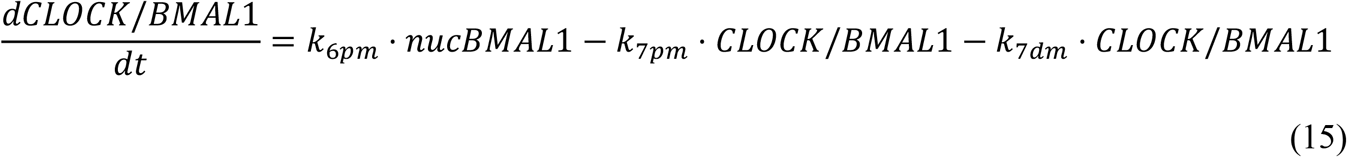

##### 1.1.3. The HPA axis mediated ISR stressor effect on central circadian clock

The exact mechanism through which the GCN2-eIF2α-ATF4 pathways within SCN neurons are activated by ISR signals and the SCN clock’s robustness is affected by unpredictable stress perturbations remains unclear. However, since the HPA axis is a major neuroendocrine system to regulate the stress response and glucocorticoids, known as the primary effectors of the HPA axis, are believed to enable the entrainment of neurotransmitter expression in the SCN [9], one hypothesis could be that stresses affect the SCN clock through an indirect extra-SCN pathway, which in this study is assumed to be the glucocorticoids-mediated stress-transducing feedback of the HPA axis on SCN neurotransmitters. In other words, our model assumed that the neurotransmitters convey both intra-SCN coupling and extra-SCN stress information.

The stress responses of the HPA axis are initiated by the release of corticotropin releasing hormone (CRH) from the paraventricular nucleus (PVN) of the hypothalamus, which then induces the production and secretion of adrenocorticotropic hormone (ACTH) by the anterior pituitary gland, followed by the output of stress hormones, glucocorticoids (CORT), from the adrenal gland (**Equation 16-18**). The stress activities of glucocorticoids, in turn, activate the expression of neurotransmitters (*V*) (the first term in **Equation 4**), transducing the stress effect to the SCN, which consequently stimulates the activation of GCN2-eIF2α-ATF4 pathway in each cell (**Equation 1**). Parameter *k*_*fb*_ in **Equation 4** denotes the strength of the CORT-dependent feedback loop to the V activities in SCN neurons.

*HPA axis:*

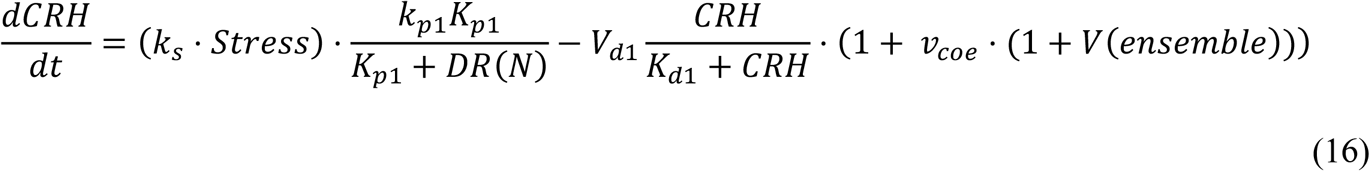

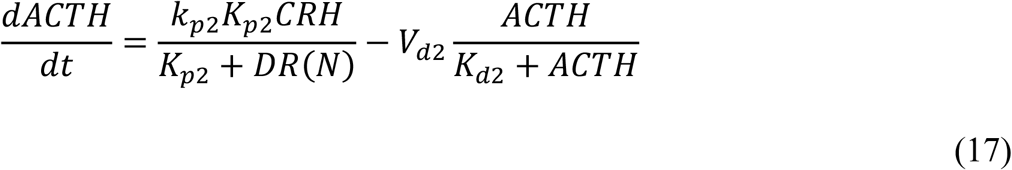

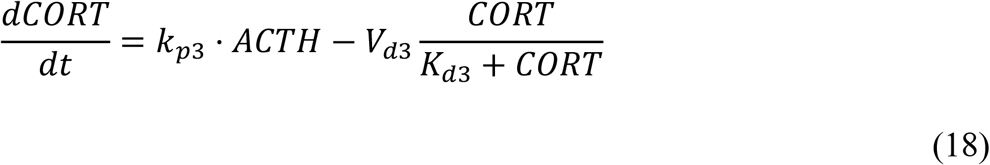

##### 1.1.4. The circadian rhythm of HPA axis and glucocorticoid receptor dynamics

Built upon our previous models [1, 2, 7], the neurotransmitter *V* released from the SCN downregulates the secretion of CRH [10] (**Equation 16**), driving the circadian rhythm of the HPA axis. Besides, a closed CORT-dependent negative feedback loop that inhibits the production of CRH and ACTH is reserved, which is essential to maintain the homeostasis and terminate the stress response of the HPA axis. Specifically, the secreted CORT binds to its receptor (*R*) (**Equation 20**) in the hypothalamus and anterior pituitary gland. Then, the formed cytoplasmic glucocorticoid-receptor complexes (*DR*) (**Equation 21**) translocate to the nucleus (*DR*(*N*)) (**Equation 22**) and exert the CORT’s negative regulation on the activities of CRH (**Equation 16**) and ACTH (**Equation 17**), as well as the receptor gene’s transcription (*R*_*mRNA*_) (**Equation 19**).

*Glucocorticoid receptor dynamics in the HPA axis:*

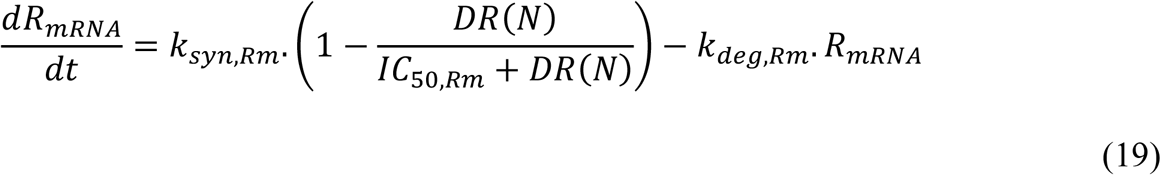

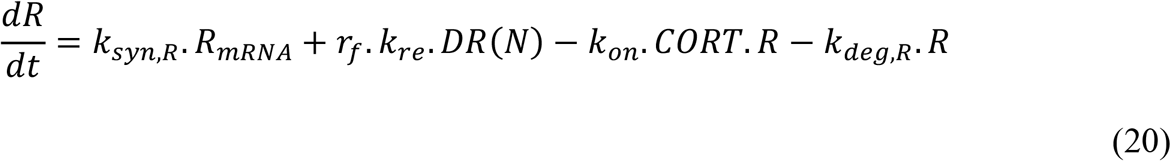

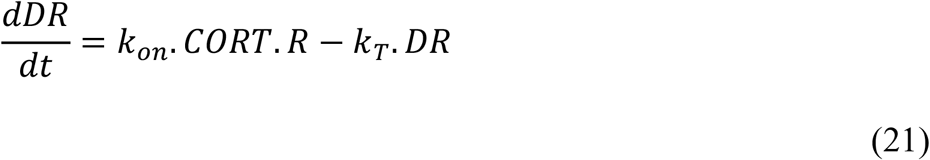

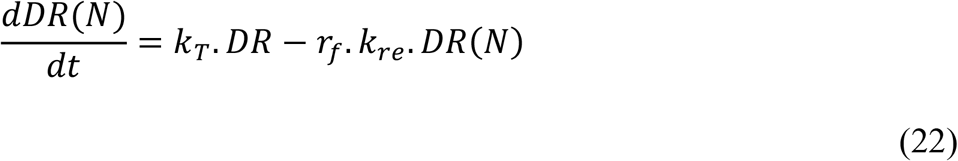

## Methods

### 1.2.1. Quantification of clock resistance to photic perturbation

A “jet lag” protocol is believed to be a useful and quantifiable evaluation tool of the SCN clock’s capability to have its unremitting oscillation resistant to an unexpected large phase shift in the external light/dark (LD) cycle, in which the SCN’s robustness against this photic perturbation is assessed by the speed of resynchronization of SCN neurons upon exposure to the sudden advance or delay of the LD cycle [11]. In this study, a 6-h advance of the 12h/12h LD cycle was introduced on the 21st day for both entrained WT and GCN2 KO individuals. An individual that has a less robust SCN clock is expected to complete the re-entrainment of the ensemble of/ the resynchronization of component neuronal oscillators faster (i.e., in fewer days).

### 1.2.2. Assessment of clock entrainment behavior

The phase response curve (PRC) and its area under curve (AUC) were used to evaluate SCN clock’s differential responses to photic and non-photic (stress) entrainers. To produce the photic stimulated PRC, a 3 h-lasting light pulse with 0.5^*^nominal light intensity was imposed to the system in constant darkness (DD) every hour and corresponding maximum ensemble phase shifts of SCN neuronal oscillators were recorded. For the non-photic stimulated PRC, a 3 h-lasting stress pulse with 5^*^nominal stress intensity was introduced to the same system in the DD environment at same time intervals, and corresponding data of the representative output were collected.

### 1.2.3. Sensitivity analysis of model response to parameters

A local sensitivity analysis approach [12] was utilized to evaluate the impact of perturbations in parameters that are associated with the newly introduced ISR (GCN2-eIF2α-ATF4) pathway and its mediated intra-SCN coupling mechanism on the completion time of the jetlag-induced re-entrainment of the SCN clock. The investigated parameters were varied by ±20%, ±50%, and ±100%, and one at a time with other parameters fixed. As the relative sensitivity indices (*SI*_*rel*_) for a varied parameter (*p*_*j*_), relative jetlag-induced ensemble re-entrainment rate (completion time) was determined as the ratio of the relative change of the state variable (*y*_*i*_(*p*_*j*_)), jetlag-induced ensemble re-entrainment rate, to the relative change of the parameter value (**Equation 23**) [13-15], where *i* represents different variation strategies employed to the parameter *p*_*j*_ and the reference/ baseline values for computing the relative changes are corresponding nominal values. Larger values of *SI*_*rel*(_*p*_*j*)_ over different parameter variations (*i* for the same *p*_*j*_) reflect a more influential parameter that significantly affects the jetlag-induced resynchronization speed of SCN neuronal oscillators, or said, the robustness of SCN clock.

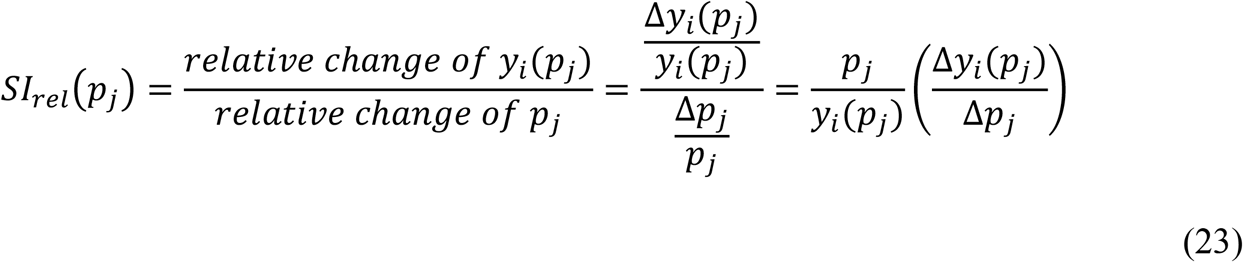

### 1.2.4. *In silico* implementation of individual variability

By hypothesizing that the stress effect on the resistance of circadian timing system, represented by CORT amplitude, is related to the functioning of intra-SCN ISR pathway, the strength of extra-SCN tissue feedback, and the perceived stress level of the system, three respectively representative parameters, *GCN*2(*T*), *k*_*fb*_, and *k*_*s*_, were sampled using Sobol algorithm to capture the individualization in these physiological properties. The resulting virtual population was further screened with the criterion that simulated CORT oscillations peak within a ±2 h window of the standard CORT peaking time (ZT12), to ensure the investigated individuals have (similar) homeostatic CORT rhythms.

### 2. Supplementary Tables

**Table S1.**
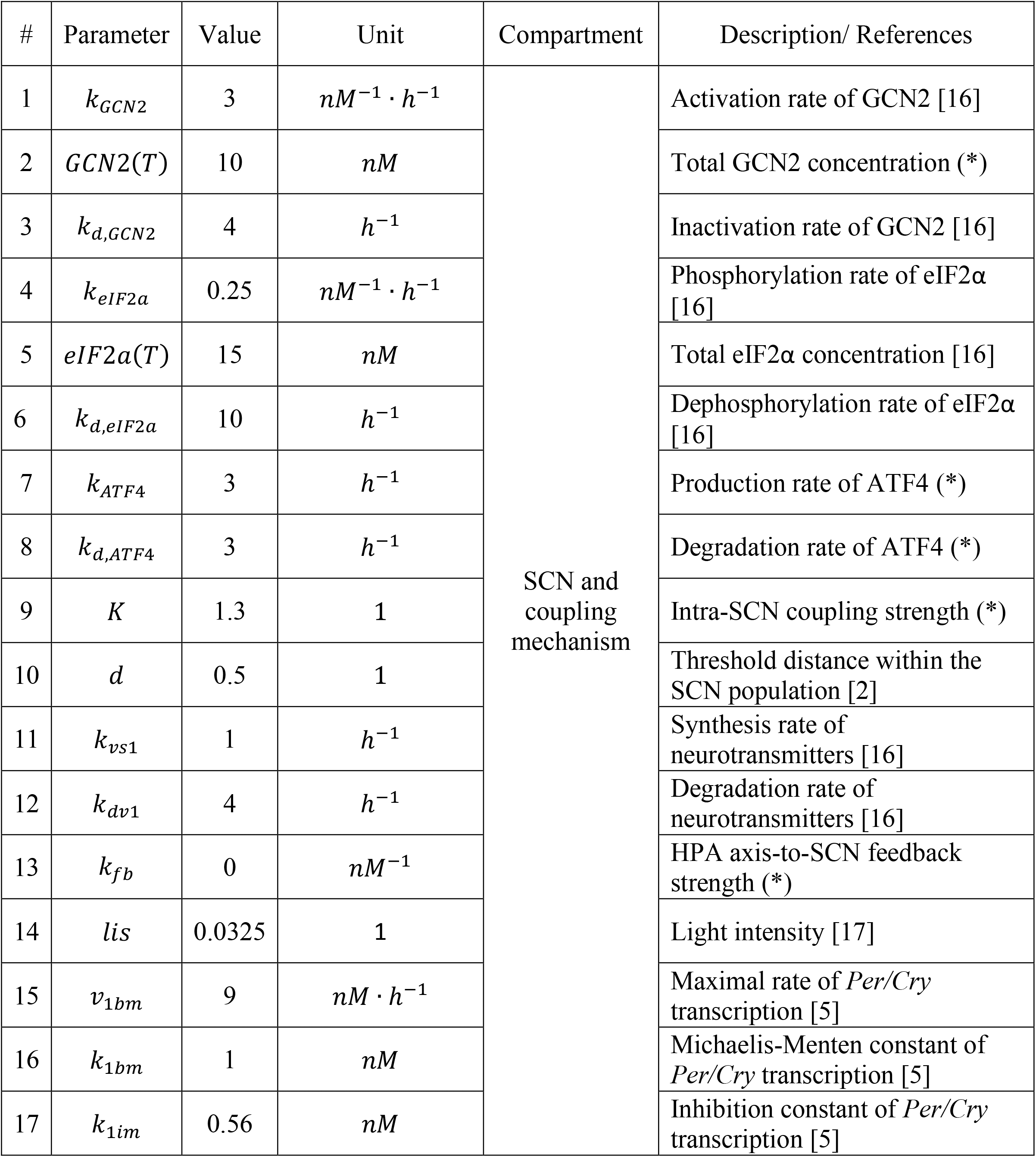
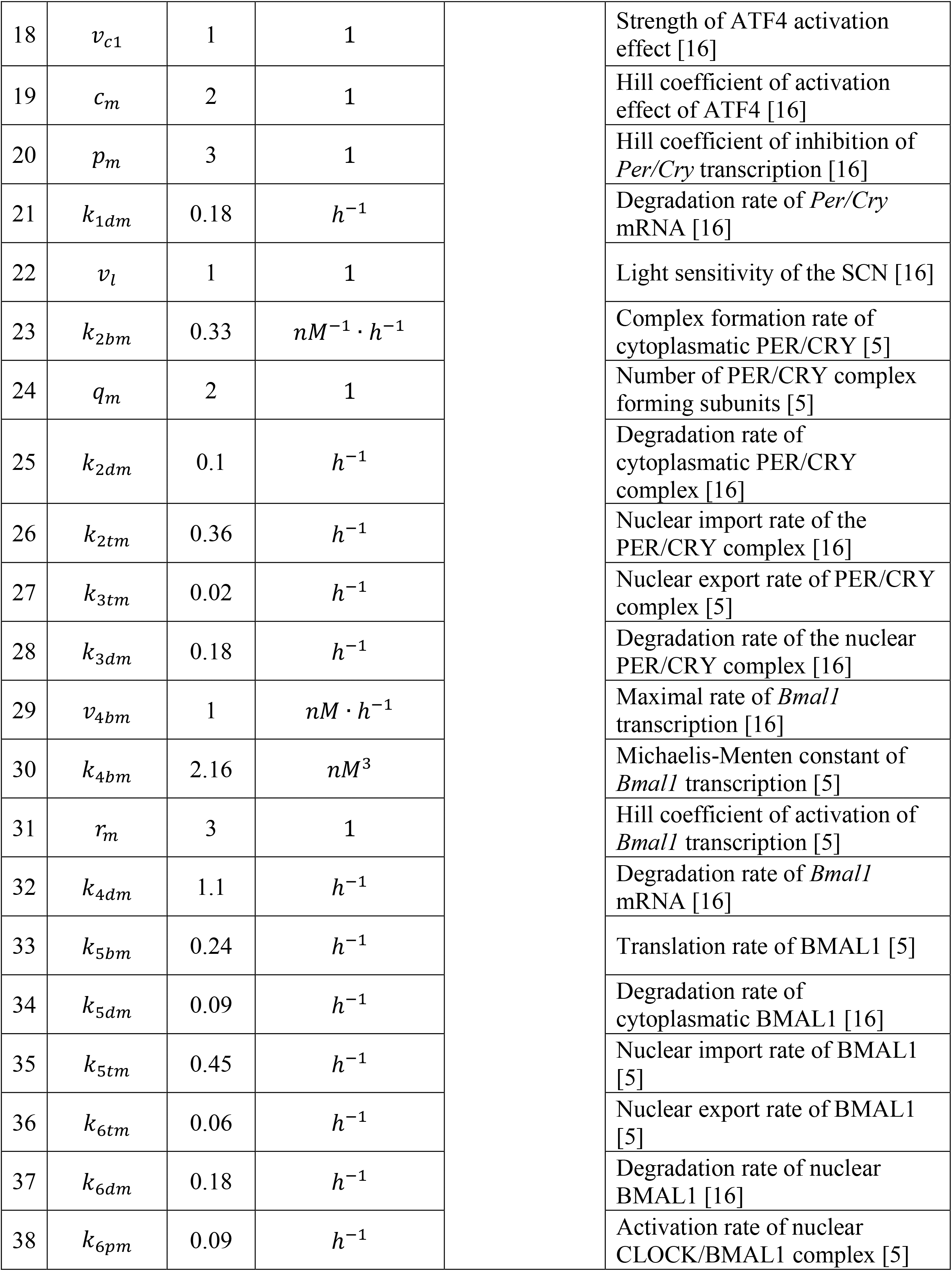
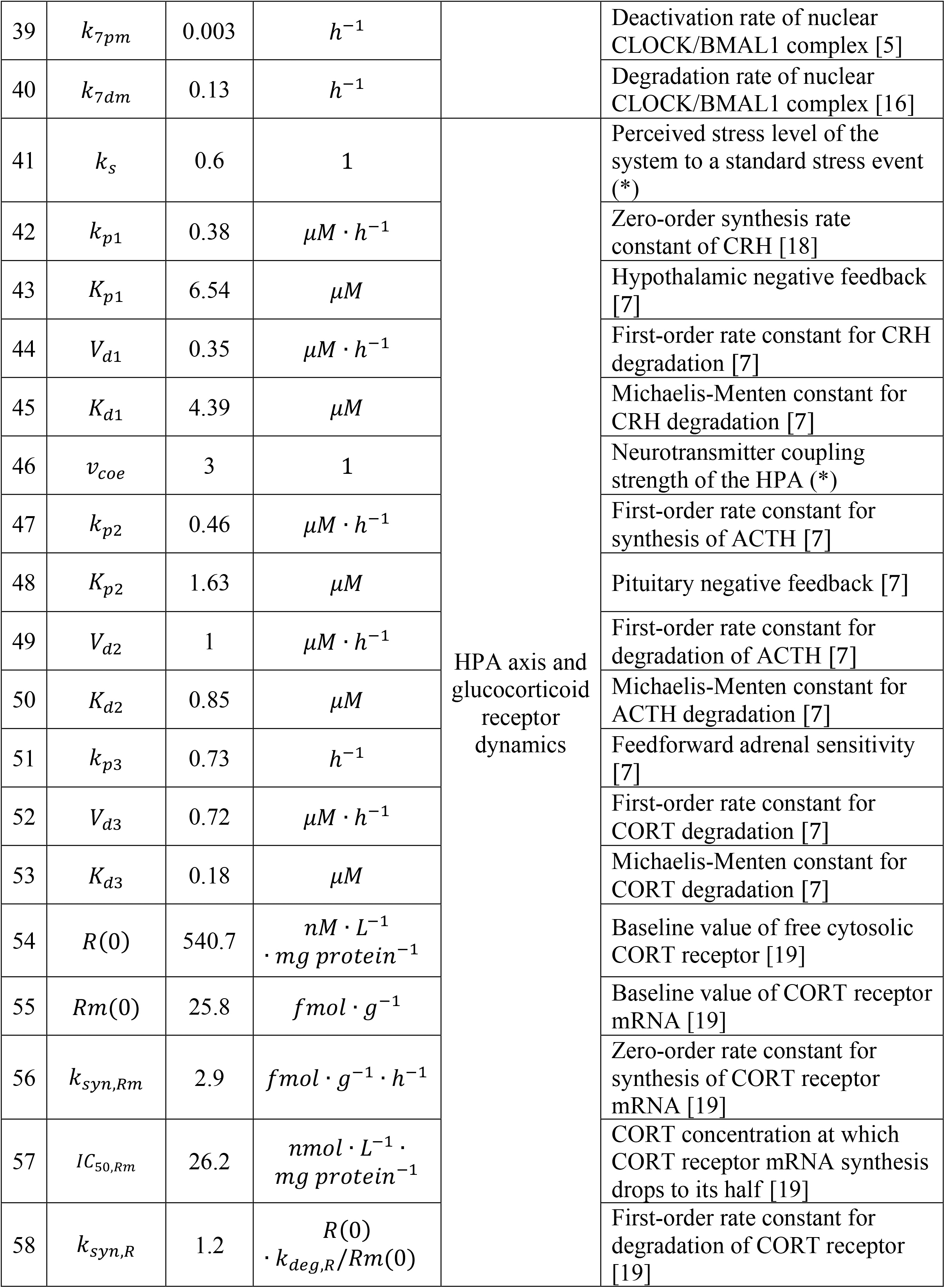
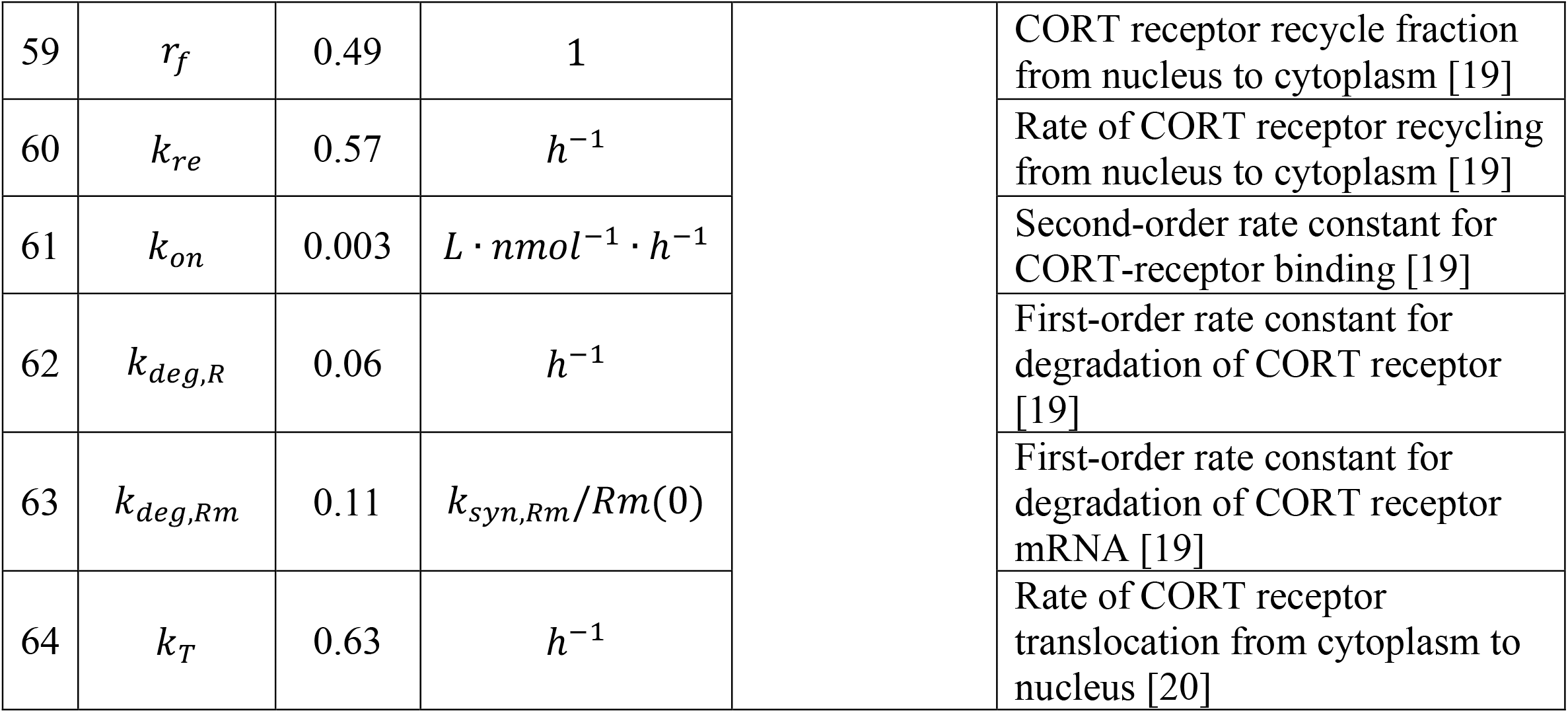
Nominal values of model parameters and their sources. (^*^) denotes the estimated parameters in this study.

